# Evolution of predators and prey kills Turing patterns

**DOI:** 10.1101/2024.03.28.587143

**Authors:** Vit Piskovsky

**Affiliations:** Mathematical Institute, University of Oxford, Woodstock Road, Oxford OX2 6GG, United Kingdom

**Keywords:** predator-prey dynamics, Turing patterns, evolution of motility, plankton patchiness, cooperation

## Abstract

The spatiotemporal patterns of predators and their prey play a pivotal role in ecology and ecological interactions can drive their formation at fine scales (1). While motility can explain the emergence of such predator-prey patterns (2–14) via the Turing mechanism (15), the predicted Turing patterns do not exhibit temporal changes that are common in experiments (16–24) and nature (25–31). Moreover, the Turing mechanism treats motility as fixed, even though predators and prey adjust their motility in response to each other (32–37) and their interactions influence their evolution (38–47). Using adaptive dynamics (48), I prove that the evolution of motility prevents the formation of Turing patterns and promotes the formation of dynamic patterns, such as predator-prey waves (28, 49–54). The resulting predator-prey cycles are shown to be induced by heterogeneous motility, which extends the emergence of predator-prey cycles beyond regimes predicted by Lotka-Volterra (55) or Rosenzweig-MacArthur (56) models. This work unites models for predator-prey spatiotemporal patterns (2–14) and evolution of motility (57–64) to explain how dynamic spatiotemporal patterns of co-evolving predators and prey emerge and persist. The novel mathematical theory is general and extends to other ecological situations, such as ecological public goods games (65).

**Significance Statement:** The spatio-temporal patterns of predators and their prey play a key role in ecology and are crucial for their conservation. Yet, even at fine scales, such patterns are often complex and exhibit spatial and temporal heterogeneity. While simple mathematical models often predict static spatial patterns (Turing patterns), I show that such patterns of predators and prey are unstable if their motility can evolve. In particular, I suggest that the evolution of motility can give rise to complex spatio-temporal patterns of predators and prey, such as predator-prey waves. Moreover, the mathematical results can be generalised to other contexts, providing novel insights into the evolution of cooperation.

---

The formation and maintenance of spatial and temporal patterns and their impact on the dynamics of populations and ecosystems are central themes in ecology (1). At different scales, patterns are governed by different mechanisms (1). For example, oceanic flow and other abiotic factors drive plankton patchiness at coarse scales, while its distribution at fine scales can emerge from biotic interactions between various constituent species (66–77). Zooplankton self-aggregates on the finest scale into swarms (66–68) to improve its predation on phytoplankton (33). Similarly, biotic interactions between acarine predators and prey can promote their spatial patterns (17, 18). Interestingly, such patterns are often dynamic. The acarine densities oscillate in the experiments (17, 18), the oceanic plankton exhibits predator-prey cycles at fine oceanic scales (31), and such predator-prey cycles are persistent in experiments with freshwater plankton (24).

While the spatiotemporal patterns of predators and prey are often dynamic in both experiments (16–24) and nature (25–31), many mathematical models either exclude space from the analysis (55, 56) or predict static spatial patterns (2– 14). The prediction of static patterns often relies on the Turing mechanism (15): random motility of predators and prey can destabilise their spatially uniform distribution. Even though the Turing mechanism is universal and can explain the formation of chemical patterns (78), patterned animal coats (79), patterns in bacterial biofilms (80) or vegetation patches in arid environments (81), it relies on a crucial assumption that the motility of interacting agents is fixed. While this assumption is reasonable if motility is passive, its application to actively moving agents deserves further justification. Predators and prey are known to adjust their motility in response to each other (32–37), and their traits including motility can evolve in response to their interactions (38–47). Although we understand how random motility leads to the formation of static spatial patterns (2–14), we have a little understanding of how such Turing patterns affect the evolution of motility and, in turn, how evolution affects Turing patterns. If the motility of predators or prey can evolve, are the Turing patterns stable on evolutionary time scales?

The evolution of motility has been modelled extensively for competing individuals of the same species (57–64), with a key prediction: slower random motility is favoured in environments with a static heterogeneous fitness landscape (57). In particular, random motility displaces the population from its fitness peaks and decreases its total fitness. The evolution of motility has also been explored in predator-prey systems, with a separate focus on predator motility (42) or prey motility (43), or with a focus on coarse scales, where the environmental heterogeneity is the primary driver of evolution (44–47). At coarse scales, motility evolves to balance the static spatial patterns introduced by variable resource quality, a prediction known as the ideal free distribution (44, 45). However, at fine scales where resource quality is relatively homogeneous, the evolution of motility in response to spatiotemporal patterns needs to be revisited.

In this work, I will explore the spatiotemporal patterns of predators and their prey at fine scales via reaction-diffusion modelling (2–14) and examine the evolution of their motility via the adaptive dynamics framework (48). In particular, I will answer the question whether the static Turing patterns, commonly predicted by many mathematical models (2–14), are evolutionarily stable.

## Results

### Model overview

To answer this question, I will model the interplay between the ecology of predators and their prey and the evolution of their motility. To commence, I construct the ecological model and recapitulate the key argument that random motility can generate static predator-prey patterns via a Turing mechanism (2–14). The ecology of predators (population *i* = *I*) and their prey (population *i* = *A*) can be described by their fecundity *f*_*i*_(*n*_A_, *n*_I_), that is by the net reproductive rate at different population densities *n*_*i*_. Generally, the fecundity of predators *f*_I_ increases with the density of prey *n*_A_ and the fecundity of prey *f*_A_ decreases with the density of predators *n*_I_ (Fig. 1a). To illustrate the general theory of this paper, I consider well-established fecundity functions (5, 6)

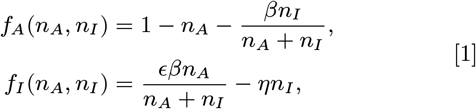

where *β* is the predation rate, *ϵ* is the conversion efficiency of consumed prey into offspring and *η* is the per capita death rate of predators. If the predators and prey are well-mixed, the fecundity functions determine the temporal dynamics of population densities *n*_*i*_(*t*), given by

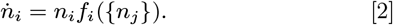

**Fig. 1.**
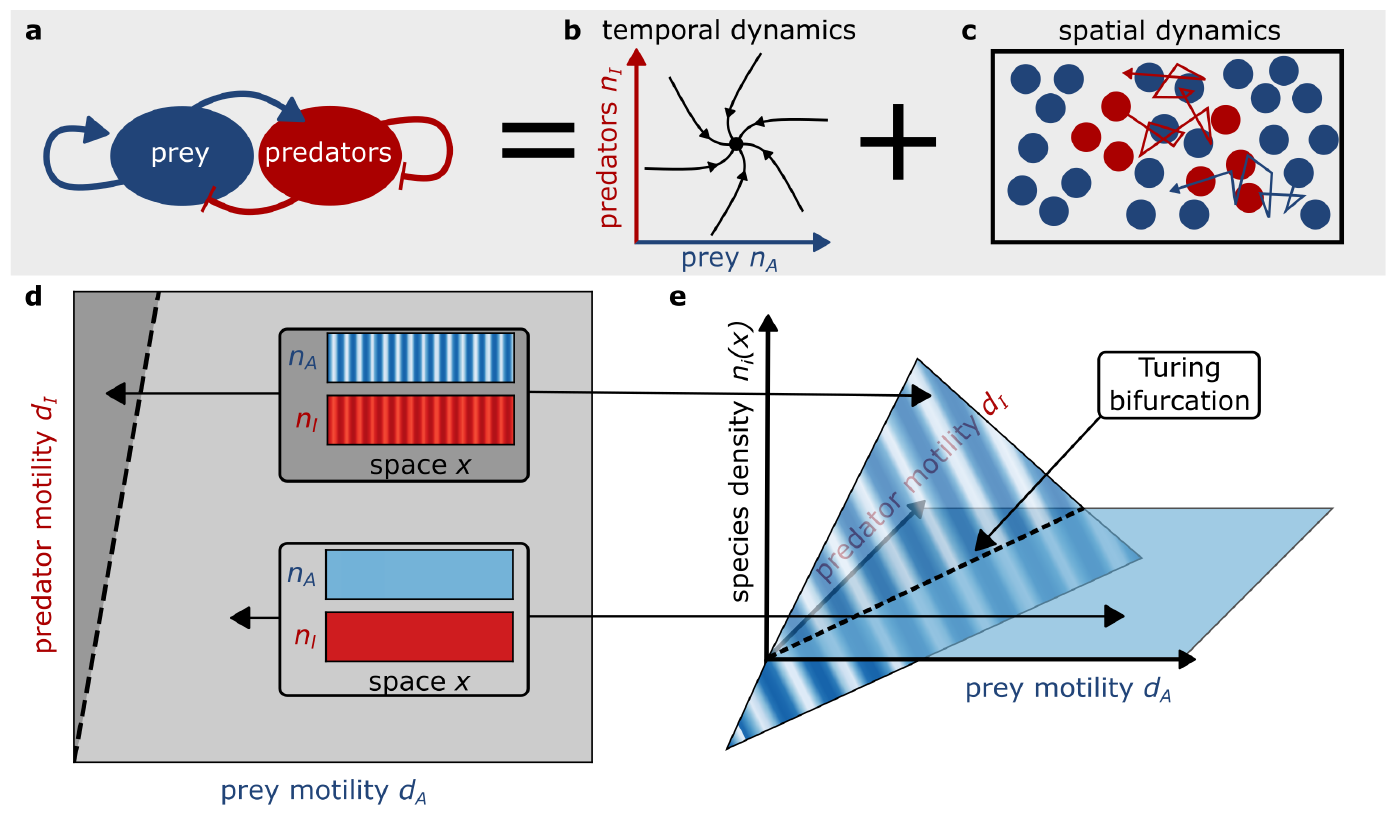
Random motility of predators and prey induces Turing patterns. (a-c) Model description. Predators (red) and prey (blue) engage in a +*/−* ecological interaction. It is assumed that the temporal dynamics admits a stable steady state (b) and that individuals of both species move randomly in space (c). (d) Pattern formation. The stable steady state of the temporal dynamics corresponds to a spatially homogeneous distribution of predators and prey, which is often stable to spatial perturbations (light grey). However, when predators are sufficiently more motile than their prey, spatial fluctuations in population densities generate Turing patterns of predators and prey (dark grey). (e) Schema of bifurcation. The Turing patterns emerge via a Turing bifurcation (dashed line): as motility changes through the bifurcation, the homogeneous steady state collides with the Turing pattern and exchanges its stability.

In this paper, I assume that the temporal population dynamics has a positive and stable steady state 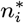 (Fig. 1b). When random motility of both species is incorporated into the model (Fig. 1c), the dynamics of the spatiotemporal population densities *n*_*i*_(*t, x*) is given by

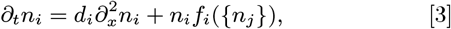

where *d*_*i*_ is the motility of species *i*. For simplicity, I assume that both species move in a one-dimensional domain of a large length *L* and cannot cross its boundary. The predator-prey dynamics in Eq. (3) admits a spatially homogeneous steady state 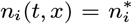 (Fig. 1d, Lemma 1 in Supporting Information Text). While this steady state is stable to spatially homogeneous perturbations in population densities, heterogeneous fluctuations in predator-prey densities can grow and introduce stable static patterns (Fig. 1d, Lemma 3 in Supporting Information Text). These static patterns are known as the Turing patterns.

#### Theorem 1

*The spatially homogeneous steady state* 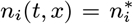 *of* Eq. (3) *is unstable to spatial perturbations (i*.*e*., *admits a Turing instability) precisely when*

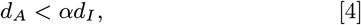

*for some α* ∈ [0, 1] *that is independent of the motility d*_*i*_ *of each species i. (See Supporting Information Text for proof*.)

Importantly, the Turing patterns form only if predators are more motile than their prey, which allows the prey to grow locally but suppresses its growth globally (Fig. 1d). At an abstract level, the spatially homogeneous steady state exchanges the stability with the Turing pattern when the motility of predators or prey passes through the line *d*_A_ = *αd*_I_, known as the Turing bifurcation (Fig. 1e). Therefore, the Turing patterns form only if *d*_A_ *< αd*_I_. This condition on motility poses an important question when motility can evolve. Does motility evolve to satisfy this necessary condition for Turing patterns?

To explore the evolution of motility, the spatio-temporal model in Eq. (3) must be generalised to incorporate multiple motility phenotypes. This can be achieved by introducing the population density 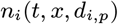 of species *i* with phenotype *p* and motility strategy 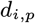 and by generalising the population dynamics in Eq. (3) to

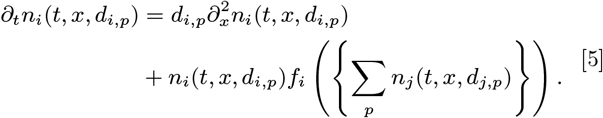

In particular, if we restrict to populations of a single motility strategy 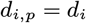, the population dynamics of the generalised model in Eq. (5) reduces back to the original model in Eq. (3) (Lemma 5 in Supporting Information Text). To study whether motility can evolve to support Turing patterns, I consider initial populations with a single motility strategy that supports such patterns, *d*_A_ *< αd*_I_, and test the hypothesis that such motility strategies are evolutionarily stable. With this aim, I model the population dynamics by Eq. (5) and the evolutionary dynamics by introducing and removing phenotypes of population *i* at a stochastic rate *µ*_*i*_. Firstly, phenotypes whose density falls below a small fixed threshold are considered extinct and are removed from the population (natural selection). Secondly, mutant phenotypes with motility strategies 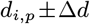 are introduced at low densities, where Δ*d*_*i*_ is a small deviation in motility around surviving phenotypes *p* of species *i* with motility strategies 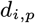 (mutations).

### Turing patterns negatively impact species fitness and motility

To understand how static spatial patterns affect the evolution of motility, I simulate the model for the predator-prey system (Movie S1) with a nearly homogeneous initial distribution of predators and prey (Fig. 2a). After a while, the motile populations form Turing patterns and predators concentrate in regions of high prey density (Fig. 2b). Surprisingly, such patterns trigger a decline in prey motility, which changes the spatial pattern (Fig. 2c). In response, predators also evolve decreased motility (Fig. 2d). Since motility allows prey to escape predators and predators to hunt prey, it is counter-intuitive that motility should decline. What is the reason behind the decline in motility of predators and their prey in this situation?

**Fig. 2.**
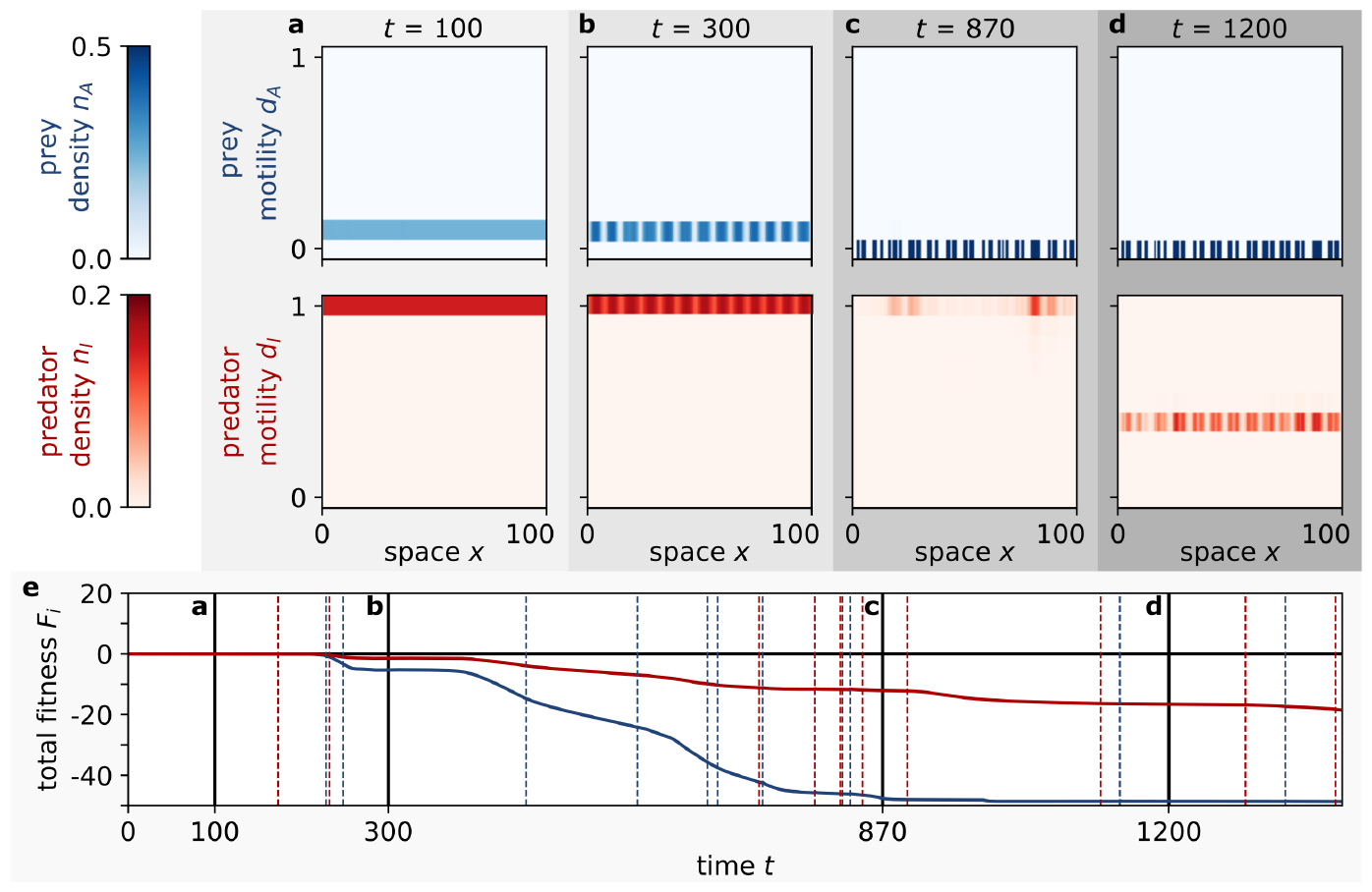
Turing patterns negatively impact species fitness and motility. (a-d) Snapshots. Prey (blue) and predators (red) have phenotypes of different motility. Initially, both populations have a single motility phenotype and are nearly spatially homogeneous (a). For appropriate initial motility, predators and prey self-organise to form Turing patterns (b). At a stochastic rate *µ*_*i*_, the phenotypes of species *i* are updated: phenotypes with a small density are considered extinct and removed (natural selection), while new phenotypes with motility perturbed around the motility of surviving phenotypes are introduced at low densities (mutations). Consequently, the prey decreases its motility (c) and so do the predators (d). In turn, changes in motility trigger changes in the Turing pattern. (e) Evolution of total fitness. The total fitness of predators (red) and their prey (blue) is defined by the total fecundity of each species. When Turing patterns form, the total fitness decreases. Vertical black lines indicate the time of snapshots in panels (a-d), horizontal black line corresponds to the optimal total fitness, and dashed lines correspond to the times when a new motility strategy emerges in prey (blue) or predators (red).

The key driver of the evolutionary dynamics is the fitness landscape experienced by each population. Crucially, when Turing patterns form, the total fitness defined by the total fecundity

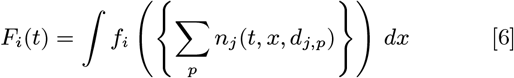

decreases for both prey and their predators (Fig. 2e).

#### Theorem 2

*If* 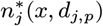 *is a steady state of the eco-evolutionary model in* Eq. (5) *and* 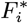 *the corresponding total fitness of species i, then the total fitness is bounded above by* 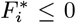 *and attains the upper bound precisely when all phenotypes p satisfy* 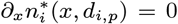, *i*.*e*., *there are no static patterns. (See Supporting Information Text for proof*.)

Therefore, once a Turing pattern forms, the total fitness of each population is decreased. The decrease in the total fitness of prey can be explained by the diffusion of prey into regions with elevated predator densities and decreased prey fecundity *f*_A_ *<* 0. Correspondingly, the total fitness of predators is decreased due to their diffusion into regions with a lack of prey and low predator fecundity *f*_I_ *<* 0. This suggests that populations with smaller motility can better utilize the local fitness peaks and are naturally selected. For example, less motile prey is less likely to migrate into regions where predators are concentrated. However, once the less motile phenotype of prey reaches fixation, the spatial pattern changes. Interestingly, while this shift in prey motility decreases the total fitness of predators, the outcome is far more devastating for the prey itself (Fig. 2e), whose total fitness is diminished even more due to increased spatial fluctuations of both populations in the new pattern. Having lost its motility completely, the prey can only wait for the predators to lose their motility as well.

This evolutionary race can be understood even better by studying the evolutionary trajectory of the motility strategies (Fig. 3a). As prey starts with a smaller initial motility than predators (Fig. 3a) and smaller total fitness induced by the pattern (Fig. 2e), prey loses its motility completely before the predators do. However, as the predators are still motile enough to form Turing patterns, the patterns prevail and predators experience a decreased total fitness, to which they respond by decreasing their motility as well. Crucially, this evolutionary dynamics can be understood analytically through the lens of the adaptive dynamics framework.

**Fig. 3.**
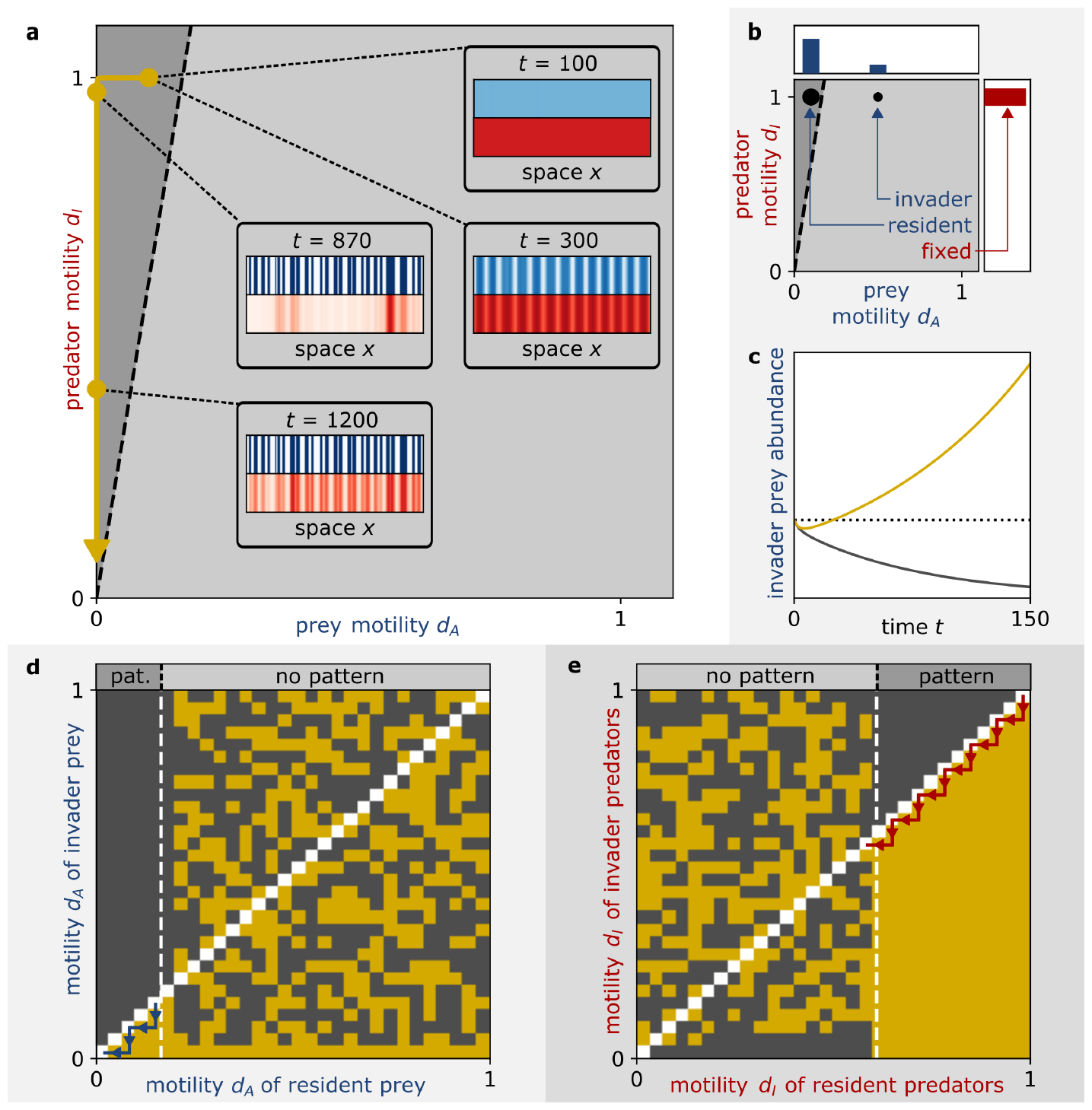
Evolution selects phenotypes of smaller motility when predators and prey form Turing patterns. (a) Evolution of motility. The evolution is indicated by the trajectory of mean predator and prey motility (golden line). The initial motility is compatible with Turing patterns (dark grey). Once patterns form, prey with smaller motility is naturally selected, followed by predators of smaller motility being selected. This leads to serial changes in the spatial patterns of predators (red) and their prey (blue), illustrated by snapshots of the total spatial density. (b-c) Invasibility analysis. The predator motility is fixed and prey with resident motility has reached a stable ecological equilibrium with predators (b). Subsequently, prey with invader motility is introduced into the system at low density (b) and the population dynamics is simulated until time 1*/µ*_A_, where *µ*_A_ is the prey mutation rate (c). If the prey with invader motility increases in abundance above its initial density (dotted line), the invader phenotype can invade the resident phenotype (yellow), otherwise not (grey) (c). (d) Pairwise invasibility plot for prey motility. The predator motility is fixed (*d*_I_ = 1) and the prey motility (*d*_A_) is varied. The white dashed line denotes the threshold for the emergence of Turing patterns in the resident population (*d*_A_ *< αd*_I_). If Turing patterns exist, smaller motility always invades (blue arrows). When Turing patterns are absent, the pairwise invasibility analysis is inconclusive and evolution is neutral. (e) Pairwise invasibility plot for predator motility. Same as (d), but the prey motility is fixed (*d*_A_ = 0.1) and the predator motility (*d*_I_) is varied. Again, if Turing patterns exist, predators with smaller motility invade (red arrows).

This framework explores phenotypic evolution by studying whether a new invader strategy can invade a population dominated by a resident strategy (48). In particular, to study how prey motility evolves, the predator motility is fixed and prey with resident motility is assumed to have reached a stable spatial equilibrium with predators. Next, an invader prey phenotype with invader motility is introduced into the prey population at low density (Fig. 3b) and both species are simulated according to the Eq. (5) until the expected time of the next mutation 1*/µ*_A_. If the invader phenotype increases in density, then the invader phenotype can invade the resident phenotype, otherwise not (Fig. 3c). By performing this invasibility analysis for prey with all pairs of resident and invader motility, the evolution of prey motility can be assessed with the same key result (Fig. 3d): when static spatial patterns form, prey with smaller motility always invades prey with higher motility. In contrast, when such patterns do not form, the fitness of prey is independent of motility, and therefore, the evolution is neutral. In particular, the abundance of the invader phenotype changes only marginally and linear pairwise invasibility analysis is not conclusive (see Methods). Furthermore, reversing the role of predators and prey in this analysis, it can be concluded that the random motility of predators is selected against when Turing patterns form (Fig. 3d). Importantly, this result can be proved analytically, as summarised for both predators and prey in the main result of this paper.

#### Theorem 3

*Assume that* 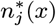 *is a positive, stable and spatially inhomogeneous steady state of* Eq. (3) *(i*.*e*., *a Turing pattern) and consider solutions of* Eq. (5)

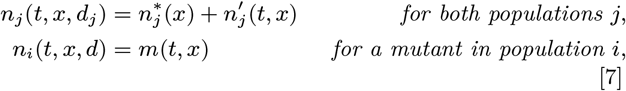

*with a mutant phenotype of species i with motility d. Then, the solution m* = 0, 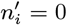 *is locally asymptotically stable (resp. unstable) if d > d*_*i*_ *(resp. d < d*_*i*_*). In particular, the steady state of* Eq. (5)

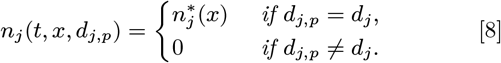

*is locally asymptotically unstable whenever there exists a phenotype p of one of the populations i with motility* 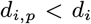 *(See Supporting Information Text for proof*.)

In summary, the motility cannot evolve to satisfy the condition for Turing patterns in Theorem 1 because phenotypes with smaller motility can always invade populations that form such patterns. Consequently, the motility of both species evolves to vanish. Put differently, motility strategies that lead to the formation of Turing patterns are not evolutionarily stable. However, what motility strategies are evolutionarily stable and attract the evolutionary dynamics? And, how do these strategies affect the spatio-temporal patterns of predators and their prey?

### Evolution of motility can introduce predator-prey waves

When predators and prey form Turing patterns, their motility evolves to vanish. However, once motility is lost, the evolution is not finished (Movie S1). Instead, the motility of predators and their prey increases and stabilises at evolutionarily stable strategies (Fig. 4a). Interestingly, the evolutionarily stable populations harbour multiple phenotypes and exhibit predatorprey waves (Fig. 4b,c). Predators hunt escaping prey, total fitness oscillates around the optimal value (Fig. 4f) and the population size exhibits typical predator-prey cycles (Fig. 4e). Surprisingly, if the populations consisted of a single phenotype given by the average motility, such dynamic spatiotemporal patterns could not form. Phenotypic heterogeneity is necessary for the formation of predator-prey waves in the presence of stabilising temporal dynamics (Lemma 2 in Supporting Information Text), even if some phenotypes are only present at very low densities (Fig. 4a-c, Theorem 4 in the Supporting Information Text). The phenotypic heterogeneity is also sufficient. If prey harbours a single motility phenotype, two predator phenotypes, slow and fast, can induce predator-prey waves (Fig. S1).

**Fig. 4.**
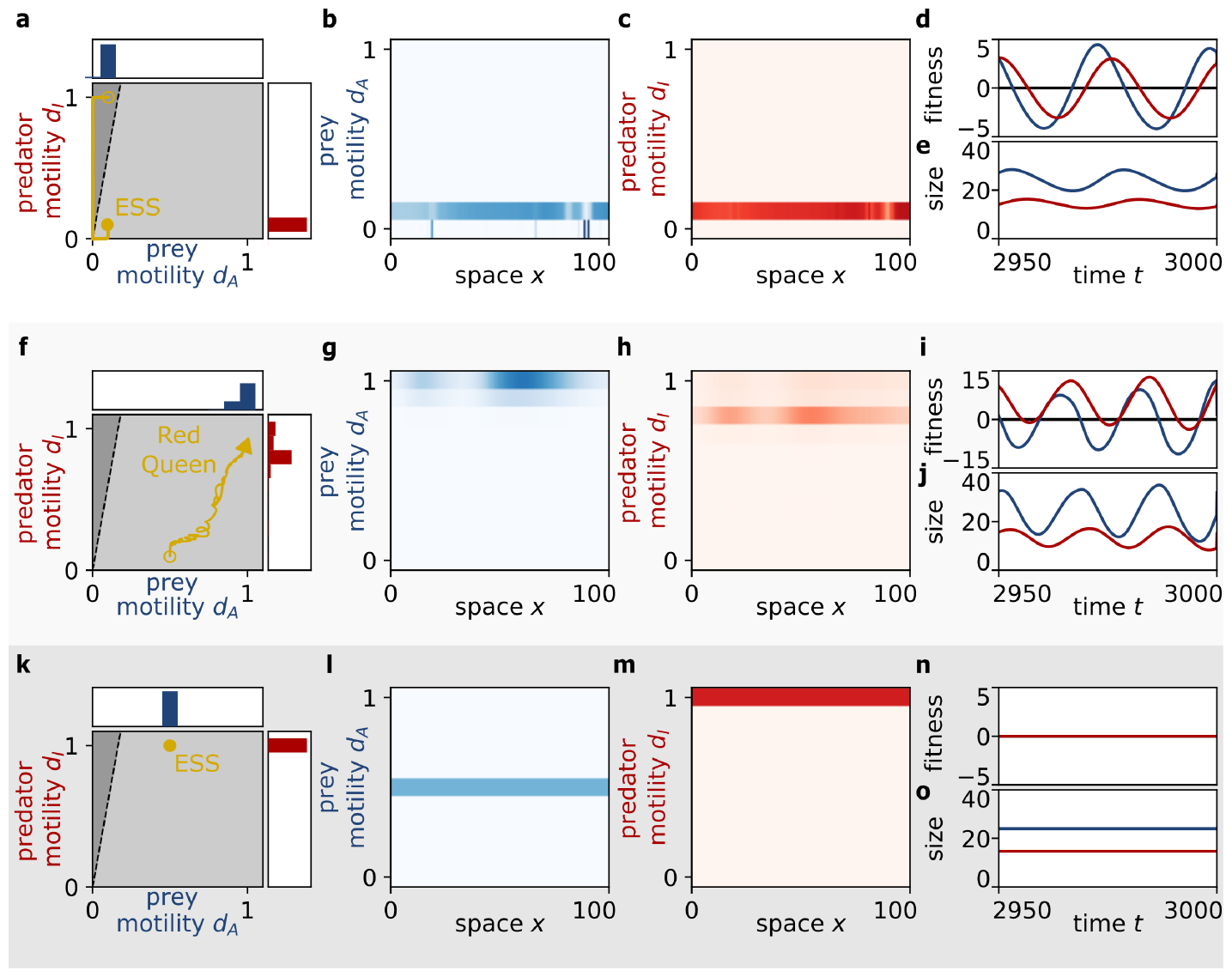
Evolution of motility can introduce predator-prey waves. (a-d) Stationary evolution, dynamic pattern. The motility of predators (red) and prey (blue) reaches an evolutionarily stable strategy (ESS, golden line). At this evolutionary equilibrium, spatio-temporal waves of prey (b) are followed by phase-shifted waves of predators (c). The total fitness of both species oscillates around its optimum (d) and the total population size (e) exhibits predator-prey cycles. (f-j) Dynamic evolution, dynamic pattern. The predators and prey evolve increasing motility (f), leading to predator-prey waves (g,h), and oscillations in total fitness (i) and total population size (j). (k-o) Stationary evolution, no pattern. The motility stays at its original value (k), and the prey (l) and their predators (m) are homogeneously distributed. The total fitness stays at the optimal value (n) and the population size does not change (o).

The emergence of predator-prey waves is relatively robust. If initial motility is chosen in the region where patterns do not form according to Theorem 1, mutations often introduce sufficient phenotypic heterogeneity for predator-prey waves to form (Fig. 4f-j, Movie S2). While the simulations exhibit the same spatio-temporal patterns, the evolution differs from the previous case. Instead of reaching an evolutionarily stable strategy, the prey increases their motility to escape their predators, forcing the predators to increase their motility to catch the faster prey, and so on (Fig. 4f). The predators and their prey are caught in an arms race, with both species evolving faster phenotypes to survive against their opponent. This evolutionary paradigm is often known as the Red Queen hypothesis (82, 83), which proposes that two opponents must constantly adapt to each other to survive. Moreover, as the predators and their prey keep evolving to hold the other species in check, the amplitude of the predator-prey waves keeps changing (Movie S2).

While predator-prey waves emerge for a large variety of initial motility values, the homogeneous density of predators and their prey can be both ecologically and evolutionarily stable if the initial predator motility is large compared to that of prey (Fig. 4k-o). However, even in these situations, the key result remains valid. The motility does not evolve into the region where static Turing patterns form. In summary, whether the resulting evolution is dynamic or stationary and patterns are homogeneous or dynamic, Turing patterns cannot evolve.

## Discussion

In this work, I have shown that random motility can affect the spatio-temporal patterns of predators and prey on fine scales, and *vice versa*, that the patterns can affect the evolution of their motility. While Turing patterns are common in mathematical models of predator-prey dynamics (2–14), random motility that induces such static patterns is not evolutionarily stable (Fig. 3). This key conclusion can be explained as an arms race (38, 40), where predators and prey continually respond to the evolution of their opponent by decreasing their motility (Fig. 2a-d). This result extends the standard result that motility is disadvantageous in static spatially heterogeneous fitness landscapes (57) to fitness landscapes that are dynamically shaped by the opponent. Since evolution selects less motile phenotypes in such landscapes, fitness costs commonly associated with motility (84, 85) are expected to accelerate this arms race. Interestingly, the loss of motility is faster in prey than in predators (Fig. 3a), even though the consequent changes in the pattern have negative impacts on the prey fitness (Fig. 2e). This supports the claim that prey evolve their traits faster than predators (40, 82, 86).

Eventually, predators and prey either reach evolutionarily stable motility strategies (Fig. 4a-e) or continually evolve increasing motility (Fig. 4f-j). Consistently with the Red Queen hypothesis, the two opponents will keep adapting to each other to survive (82, 83). These forms of evolution commonly lead to the emergence of predator-prey waves (Fig. 4). While fixed random motility dampens predator-prey oscillations (87) and stabilises homogeneous predator-prey distribution (Fig. 1d, Theorem 1), I show that multiple motility phenotypes can induce predator-prey oscillations (Fig. 4a-j, Theorem 4 in Supporting Information Text). Waves of predators that chase prey emerge in other models (28, 49–54) and, unlike Turing patterns, are stable under the evolution of motility (51). Indeed, predator-prey oscillations are common in natural (25–31) and experimental (16–24) settings, and my model provides an evolutionary robust mechanism for their formation. This mechanism introduces predator-prey cycles via heterogeneous motility (Fig. 4a-j, Lemma 2 and Theorem 4 in Supporting Information Text) and proves that predator-prey cycles can emerge even if the temporal dynamics in Eq. (2) is stable. The predator-prey cycles in this work can be contrasted with cycles in Lotka-Volterra (55) or Rosenzweig–MacArthur models (56) which require oscillatory temporal dynamics. Consequently, heterogeneous motility might offer a novel explanation for the predator-prey oscillations of oceanic plankton at fine scales (31) and the persistence of such oscillations in experiments with freshwater plankton (24), including irregular transitions in the oscillatory pattern (Movie S2).

The model also contributes to the field of adaptive dynamics (48, 88), extending it to the contexts of spatial movement and co-evolution of multiple species. Importantly, adaptive dynamics assumes that mutations generate small changes to the present motility strategies. However, actively moving organisms can quickly switch between motility strategies that are very different (33) and the presence of such large-effect mutations can modify the outcomes of evolution (89). However, even if mutations of large-effect are included in the modelling (see Methods), the key conclusion remains valid: motility compatible with the formation of static Turing patterns is not evolutionarily stable (Fig. S2, Movie S3).

While this conclusion has been achieved for the case of predation, the presented mathematical theory applies to arbitrarily many interacting populations that form static Turing patterns (Theorem 3 in Supporting Information Text). When two populations are considered, the theory extends to any population with a +*/*− ecological interaction, a type of interaction that is necessary for patterns to form (Lemma 4 in Supporting Information Text). For example, cooperators and defectors that produce (resp. exploit) a common resource in an ecological public goods game have been predicted to create Turing patterns (65) (see Methods). Despite the predicted emergence of Turing patterns, the corresponding motility of cooperators and defectors is not evolutionarily stable (Fig. S3 and S4, Movie S4) and cooperators and defectors evolve reduced motility. However, once motility is lost, the resulting dynamics can be different from predators and prey (Fig. S5). Unlike prey that can survive in the absence of predators, cooperators require a sufficient density for survival and a simultaneous extinction of cooperators and defectors is temporally stable (65). Therefore, when cooperators and defectors lose motility, they coexist in regions where the Turing pattern peaked and they go extinct in regions where the Turing pattern exhibited low organismic densities. Consequently, the vanishing motility is evolutionarily stable and the static spatial pattern becomes imprinted into the system (Fig. S5a-f, Movie S4). Nevertheless, these results need to be challenged as motility can explicitly affect organismic fitness, contradictory to the assumptions of this model. For example, *Paenibacilli* cooperate to produce a lubricant that allows them to move across hard surfaces (90) and *Myxococci* cooperatively create extracellular fibril matrix to swarm (91).

While this work addresses how random motility and spatiotemporal patterns of organisms affect each other, many organisms can bias their movement. For example, *E. coli* can cooperatively excrete a chemical signal that attracts other cells and leads to pattern formation (92, 93). Motility bias is also common in predator-prey systems as predators move towards high prey densities and prey moves away from predators (32). Oceanic plankton aggregates into swarms at high phytoplankton densities (33), seabirds follow high fish densities (35), whereas tadpoles move away from their dragonfly predators (34, 36), and the presence of predators can trigger the production of winged offspring in aphids (94). It has been separately suggested that biased motility promotes the formation of patterns (8, 95, 96) and the evolution of motility (60–64). This model might provide a starting point for exploring how biased motility and patterns of ecologically interacting organisms affect each other.

In summary, this work shows that the evolution of motility constrains spatio-temporal patterns of predators and prey. While mathematical models predict spatially heterogeneous patterns that can be both static (Turing patterns (2–14)) or dynamic (predator-prey waves (28, 49–54)), patterns that occur in experiments (16–24) and nature (25–31) are often dynamic. Here, I show that the static Turing patterns are unstable if organismic motility can evolve, whereas the dynamic patterns are often evolutionarily stable (51). This work unites models for the predator-prey patterns (2–14) and the evolution of motility (57–64), and extends previous models that focused separately on the evolution of predators (85) and prey (43). More generally, this work suggests that the evolution of predators and prey (38–47, 97) can have an impact on their spatio-temporal patterns at fine scales, such as on the distribution of oceanic plankton (66–77).

## Materials and Methods

### Fecundity functions

The parameters of the fecundity functions for the predator-prey system in Eq. (1) are chosen as *β* = 2, *η* = 0.62 and *ϵ* = 0.5 for the illustration of the Turing patterns. Moreover, the positive steady state 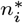 of the homogeneous system Eq. (2) can be found explicitly as 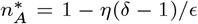 and 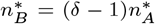 (5) and its stability can be checked by direct computation of the Jacobian *J*(*k*^2^) introduced in Section 1 of Supporting Information Text. For defectors (*I*) and cooperators (*A*), I use the fecundity functions from (65)

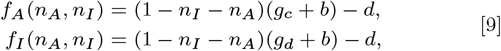

where *b* is the base-line birth rate, *d* is the per capita death rate and *g*_c,d_ are the birth rates of cooperators and defectors associated with producing and utilizing a common resource

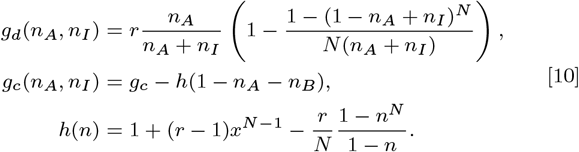

The parameters are chosen as *r* = 2.4, *N* = 8, *b* = 1 and *d* = 1.2 to illustrate the case where Turing patterns emerge. Since the positive steady state 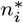 of the homogeneous system in Eq. (2) cannot be found explicitly (65), the Newton-Raphson method is used to locate it. A direct computation of the Jacobian in terms of 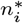 again reveals that this steady state is stable (65). Moreover, the trivial fixed point *n*_i_ = 0 is also stable in this system (65).

Finally, I note that the direct computation of the Jacobian is also used to plot the line *d*_A_ = *αd*_I_ of the Turing bifurcation in Fig. 1d, Fig. 3a,b, Fig. 4a,f,k, Fig. S2a,f,k, Fig. S4a,b and Fig. S5a,f,k. The coefficient *α* can be expressed in terms of the Jacobian, as shown in the proof of Theorem 1 (see Supporting Information Text).

### Discretization

A major component of the methods is computer simulations of the eco-evolutionary model from the main text. This model extensively uses Eq. (5) for the time-evolution of population densities 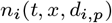. To numerically simulate Eq. (5), the independent variables *t, x* must be discretized and the discretization of motility 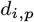 must be specified. Hereby, I introduce the appropriate discretization schemes.

Motility can be discretized by considering 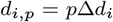, where 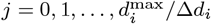. The system of partial differential equations in Eq. (5) can be reduced to ordinary differential equations via the method of lines. In particular, the space can be discretized as *x*_k_ = *k*Δ*x* and 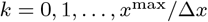 and the spatial derivative in Eq. (5) is replaced by second-order central difference

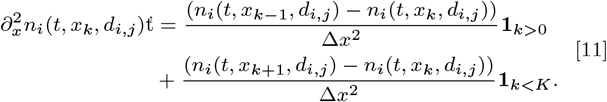

After substitution of Eq. (11) into Eq. (5), one obtains the corresponding ordinary differential equations for population densities 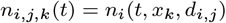 in the form

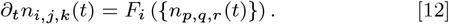

These ordinary differential equations are numerically simulated by the Euler method with time-step Δ*t*

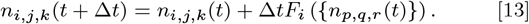

As a final remark, the spatial integral of any function *g*(*x*) is computed with the trapezoidal rule

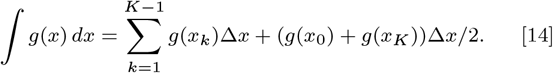

### Simulations

This section presents the algorithm for simulating the eco-evolutionary model from the main text. This algorithm describes how population densities 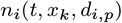 evolve in time and is inspired by the algorithm in (88). I use the discretization scheme as above with the parameters Δ*d*_A_ = Δ*d*_I_ = 0.1, 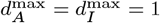, *x*^max^ = 100, Δ*t* = 0.1, Δ*x* = 0.5. The eco-evolutionary model further requires the maximal simulated time *t*_max_, the parameters that specify the initial conditions and the parameters that specify the rules for updating phenotypes. Initial conditions are specified by the size of the initial spatial perturbation *p* = 0.0001 and the initial motilities *d*_A_ = 0.1 and *d*_I_ = 1. The phenotypes are updated according to specified mutation rates for each population *µ*_A_ = *µ*_I_ = 1*/*150 and the maximal local density of introduced mutants *m* = 0.0001. The algorithm has several dynamically changing variables: the population densities *n*_i_(*t, x*_k_, *d*_i,p_), the current time *t*, the time *T* when the next mutation occurs in one of the populations and the population *M* that mutates. In the algorithm, *S, S*^′^, *S*_k_ denote numbers that are chosen uniformly at random from the interval [0, 1] and independently from any previously chosen numbers. The algorithm combines Gillespie’s algorithm for updating the phenotypic structure and the method of lines for solving the population dynamics in Eq. (5). Specifically, the algorithm consists of the following steps:

1. Initialization: Set the initial time *t* = 0. Set the initial population densities as 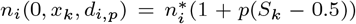 if 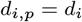, and 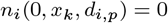 otherwise. Sample the time of the next mutation *T* = − ln *S/*(*µ*_A_ + *µ*_I_). Sample the population that mutates as *M* = *A* if *S*^′^ *< µ*_A_*/*(*µ*_A_ + *µ*_I_), and *M* = *I* otherwise.
2. Update: Repeat the steps below until *t > t*_max_.
  a. If *t > T*, update phenotypes of the population *M* :
    i. Remove phenotypes *j* with low density: If 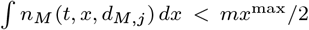 for some *j*, then set 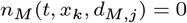 for all *k*.
    ii. Introduce mutant phenotypes at low density: set 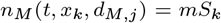 if 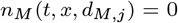 (phenotype *j* is not present) and 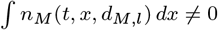 for *l* = *j* − 1 or *l* = *j* + 1 (a phenotype with nearby motility strategy is present).
    iii. Find new *T* and *M* : Sample the time of the next mutation *T* = *t* − ln *S/*(*µ*_A_ + *µ*_I_) and the population that mutates *M* = *A* if *S*^′^ *< µ*_A_*/*(*µ*_A_ + *µ*_I_), and *M* = *I* otherwise.
  b. Update densities 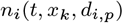 via Eq. (13).
  c. Update the time *t* → *t* + Δ*t*.

This algorithm treats the small-effect mutations, as discussed in the main text. When large-effect mutations are considered, the requirement for phenotypes with nearby motility strategies is removed from step 2(a)ii and all absent phenotypes are introduced at this step. This algorithm was used to produce Fig. 2 (*t*_max_ = 1400), Fig. 3a (*t*_max_ = 1500), Fig. 4 (*t*_max_ = 3000), Fig. S2 (*t*_max_ = 3000), Fig. S3 (*t*_max_ = 1400), Fig. S4a (*t*_max_ = 1500), Fig. S5 (*t*_max_ = 3000).

### Pairwise invasibility analysis

To produce Fig. 3c-e and Fig. S4c-e, pairwise invasibility analysis was used. For a chosen population *i*, this analysis tests whether a rare phenotype with invader motility *d*_inv_ can invade a steady state of a phenotype with resident motility *d*_res_ when the motility of the entire opponent population is fixed at *d*. Firstly, the steady state of the system without the invader phenotype is located. Both populations are initialized at 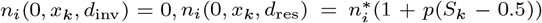 and 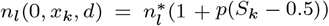 for *l* ≠ *i. S*_k_ are independent random numbers chosen uniformly from [0, 1]. The system is further evolved until the time *t* = 300, a sufficiently long time for the populations to reach a steady state. Next, the invader phenotype is introduced into the system at a low density 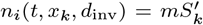, where 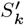 are additional independent random numbers chosen uniformly from [0, 1]. Therefore, the initial abundance of the invader phenotype is *M*_0_ = *n*_i_(*t, x, d*_inv_) *dx*. The system is evolved by additional time *t*^′^ = 1*/µ*_i_, which corresponds to the expected time when phenotypes with low abundance are removed from the system (see the algorithm above). After this time, the abundance of the invader phenotype is *M*_*t*_^′^ = ∫ *n*_*i*_(*t* + *t*^′^, *x, d*_inv_) *dx*. Since the density threshold for removing a phenotype is chosen to match the expected density of introduced mutants 𝔼*M*_0_ = *mx*^max^*/*2, the invasion is defined as successful precisely when *M*_*t*_^′^ > *M*_0_.

Indeed, Theorem 3 suggests that generically 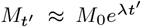 for 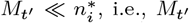, i.e., *M*_*t*_^′^ grows exponentially at rate *λ*. The rate *λ* = ln(*M*_*t*_^′^*/M*_0_)*/t*^′^ is the invasion exponent, and the criterion for successful invasion can be rewritten as *λ >* 0 (48). By following the proof of Theorem 3, it can be shown that if the steady state of Eq. (3) is spatially homogeneous, then to leading order *λ* = 0. This indicates that the pairwise invasibility analysis is degenerate and *M*_*t*_^′^ changes only marginally in the absence of patterns. Therefore, the results in Fig. 3d,e and Fig. S2d,e shall be read with caution, noting that *M*_*t*_^′^ ≈ *M*_0_ when the resident motility supports a spatially homogeneous steady state.

To produce Fig. 3d,e and Fig. S4d,e, the invasibility analysis is repeated for each pair of resident *d*_res_ and invader *d*_inv_ motility strategies that are not equal and which attain the discretized values with Δ*d*_A_ = Δ*d*_I_ = 1*/*30 and 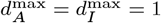.

## Code availability

All code will become available upon publication at:

https://github.com/vit-pi/MotilityEvolutionPatterns.

## ACKNOWLEDGMENTS

I thank P. Maini, K. Foster, P. Piskovska, T.J. Jewell, V. Klika, P. Haas, N.M. Oliveira and R. Van Gorder for helpful comments on this manuscript. V.P. was supported by the Mathematical Institute Scholarship.

