## Supporting Information for "Evolution of predators and prey kills Turing patterns"

**Vit Piskovsky**

**Corresponding Author Vit Piskovsky.**
****

#### This PDF file includes:

Supporting text
Figs. S1 to S5
Legends for Movies S1 to S4
SI References

#### Other supporting materials for this manuscript include the following:

Movies S1 to S4

### Supporting Information Text

The Supporting Information Text includes a systematic derivation of all analytical results:

- Section 1 introduces a general model for the spatiotemporal dynamics of interacting ecological species with random motility. The homogeneous distribution of species is identified as a steady state of the dynamics (Lemma 1) and a classification of motility-induced spatiotemporal patterns is provided.

- Section 2 explores the spatiotemporal dynamics of two species with fixed motility. While fixed motility of two species cannot induce dynamic patterns (Lemma 2), static patterns can emerge (Lemma 3) if the two species are involved in a $+/-$ ecological interaction (Lemma 4) and the predator motility is suitably larger than the prey motility (Theorem 1).

- Section 3 explores the evolution of motility if static patterns emerge. Static patterns decrease the fitness of all species that create the pattern (Theorem 2). Moreover, the static patterns correspond to steady states of the evolutionary dynamics (Lemma 5) and are evolutionarily unstable (Theorem 3).

- Section 4 explores how phenotypic heterogeneity introduced by mutations affects the spatiotemporal patterns. Heterogeneous motility can give rise to static and dynamic patterns (Theorem 4).

### 1. General spatiotemporal dynamics

The ecological interactions between species $i$ are governed by their fecundities $f_i(\{n_j\})$, where $n_j$ is the abundance of species $j$. In well-mixed systems, the temporal dynamics of these species is given by Eq. (2)

$$\dot{n}_i(t) = n_i(t) f_i(\{n_j(t)\}). \qquad \text{[S1]}$$

Since the goal is to study Turing patterns, I assume this dynamics admits a positive and stable steady state $n_i^*$. When random motility is incorporated into the model, the dynamics of the spatiotemporal population densities $n_i(t,x)$ is given by Eq. (3)

$$\partial_t n_i = d_i \partial_x^2 n_i + n_i f_i(\{n_j\}), \qquad \text{[S2]}$$

where $d_i$ is the motility of species $i$. For simplicity of presentation, I assume that both species move in a one-dimensional domain of a large length $L$ ($x \in [0, L]$) and cannot cross its boundary ($\partial_x n_i|_{x=0,L} = 0$).

**Lemma 1.** *The spatio-temporal dynamics in* Eq. (S2) *admits a spatially homogeneous steady state* $n_i(t,x) = n_i^*$.

*Proof.* Notice that Eq. (S1) implies that $f_i(\{n_j^*\}) = 0$ and $\partial_x^2 n_i^* = 0$. Therefore, $d_i \partial_x^2 n_i^* + n_i^* f_i(\{n_j^*\}) = 0$. □

The spatiotemporal stability of the steady state $n_i^*$ in Eq. (S2) can be studied by considering a complete basis of eigenfunctions for the operator $\partial_x^2$. This operator has a complete basis of eigenfunctions $\cos(kx)$, where $k = \pi n/L$ denotes a wavelength and $n \in \mathbb{Z}$. Since $L$ is assumed to be large, we can effectively assume that $k \in \mathbb{R}$. Crucially, the steady state $n_i^*$ of the spatiotemporal system in Eq. (S2) is stable precisely if it is stable for all spatial perturbations $n_i(t,x) = n_i^* + \epsilon_i(t) \cos kx$ of all wavelengths $k$. Linearising Eq. (S2) and considering $\epsilon_i(t) \sim e^{\lambda t}$, it follows that the steady state $n_i^*$ is stable precisely when the eigenvalues $\lambda(k^2)$ of the Jacobian

$$J(k^2) = \begin{pmatrix} n_1^* \partial_1 f_1(\{n_j^*\}) - d_1 k^2 & n_1^* \partial_2 f_1(\{n_j^*\}) & \dots \\ n_2^* \partial_1 f_2(\{n_j^*\}) & n_2^* \partial_2 f_2(\{n_j^*\}) - d_2 k^2 & \dots \\ \vdots & \vdots & \ddots \end{pmatrix}. \qquad \text{[S3]}$$

have negative real parts for all wavelengths $k$. When $k = 0$, the Jacobian $J(0)$ coincides with the Jacobian of the temporal system in Eq. (S1) evaluated at the steady state $n_i^*$. By assumption, the steady state $n_i^*$ of the temporal system in Eq. (S1) is stable and the Jacobian $J(0)$ has eigenvalues $\lambda(0)$ with negative real parts. Since the eigenvalues $\lambda(k^2)$ vary smoothly with $k$, the steady state $n_i^*$ is unstable precisely if some eigenvalue crosses the line $\mathrm{Re}\ \lambda(k^2) = 0$ from $\mathrm{Re}\ \lambda(k^2) < 0$ to $\mathrm{Re}\ \lambda(k^2) > 0$ as $k$ varies. Moreover, as $J(k^2)$ has real entries and the corresponding characteristic equation has real coefficients, the eigenvalues $\lambda(k^2)$ are either real or appear in complex conjugate pairs. If a real eigenvalue $\lambda(k^2)$ crosses the boundary, the resulting instability suggests the formation of static patterns, commonly known as Turing patterns. If a complex pair of eigenvalues $\lambda(k^2)$ crosses the boundary, the resulting instability suggests the formation of dynamic patterns. In particular, we can make the following definitions.

**Definition 1.** *The steady state $n_i^*$ admits a Turing instability whenever a real eigenvalue $\lambda(k^2) \in \mathbb{R}$ of $J(k^2)$ crosses the boundary $\lambda(k^2) = 0$ at some wavelength $k$. The steady state $n_i^*$ admits a Turing-Hopf instability whenever a complex eigenvalue $\lambda(k^2) \notin \mathbb{R}$ of $J(k^2)$ crosses the boundary $\lambda(k^2) \in i\mathbb{R} \setminus \{0\}$ at some wavelength $k$.*

In summary, Turing instability indicates the presence of (static) Turing patterns, while Turing-Hopf instability indicates the presence of dynamic patterns.

### 2. Spatiotemporal dynamics of two species with fixed motility

In the case of two interacting ecological species, the spatiotemporal stability of the steady state $n_i^*$ can be determined analytically. In particular, two species with fixed motility cannot exhibit dynamic patterns.

**Lemma 2.** *Two species with fixed motility cannot admit a Turing-Hopf instability.*

*Proof.* Let $J(0)$ be the Jacobian of the temporal system in Eq. (S1) evaluated at the steady state $n_i^*$. Since the steady state is stable, the eigenvalues of $J(0)$ have negative real parts, or equivalently tr $J(0) < 0$ and det $J(0) > 0$. If the system admitted a Turing-Hopf instability, then there is a wavelength $k$ such that the Jacobian

$$J(k^2) = J(0) - k^2 \begin{pmatrix} d_1 & 0 \\ 0 & d_2 \end{pmatrix}. \quad \text{[S4]}$$

has a pair of purely complex eigenvalues, that is tr $J(k^2) = 0$. However,

$$\text{tr } J(k^2) = \text{tr } J(0) - k^2(d_1 + d_2) < 0, \quad \text{[S5]}$$

leading to a contradiction. □

While dynamic patterns cannot be achieved in a two-species model, Turing patterns can emerge.

**Lemma 3.** *Two species admit a Turing instability precisely when*

$$d_1 n_2^* \partial_2 f_2(\{n_j^*\}) + d_2 n_1^* \partial_1 f_1(\{n_j^*\}) > 2\sqrt{d_1 d_2 det\ J(0)}. \quad \text{[S6]}$$

*Proof.* The steady-state $n_i^*$ admits a Turing instability precisely if the Jacobian $J(k^2)$ has a real eigenvalue that passes through the boundary $\lambda(k^2) = 0$ for some $k$. If this happens, the other eigenvalue must be real and negative. Otherwise, a complex pair of eigenvalues would contradict that a two-dimensional system has two eigenvalues and a positive eigenvalue would contradict that the sum of eigenvalues is negative, i.e., tr $J(k^2) <$ tr $J(0) < 0$. Therefore, the steady-state $n_i^*$ admits a Turing instability precisely when det $J(k^2)$ passes through the boundary det $J(k^2) = 0$ at some $k$, where

$$\text{det } J(k^2) = k^4 d_1 d_2 - k^2[d_1 n_2^* \partial_2 f_2(\{n_j^*\}) + d_2 n_1^* \partial_1 f_1(\{n_j^*\})] + \text{det } J(0). \quad \text{[S7]}$$

The function det $J(x)$ is quadratic function in $x = k^2 \geq 0$, satisfies det $J(0) > 0$ and has a positive leading coefficient $d_1 d_2 > 0$. Therefore, det $J(x)$ passes through 0 at some positive $x$ if it admits a minimum at a non-negative value $x'$

$$0 \leq \frac{d_1 n_2^* \partial_2 f_2(\{n_j^*\}) + d_2 n_1^* \partial_1 f_1(\{n_j^*\})}{2 d_1 d_2} = x'$$

and the minimal value is negative

$$0 > -\frac{[d_1 n_2^* \partial_2 f_2(\{n_j^*\}) + d_2 n_1^* \partial_1 f_1(\{n_j^*\})]^2}{4 d_1 d_2} + \text{det } J = \text{det } J(x')$$

Finally, these two conditions can be restated as a single condition in Eq. (S6). □

Importantly, the Turing patterns can only emerge if the two species exhibit a $+/-$ ecological interaction, such as predation (Fig. 1a).

**Lemma 4.** *Two species that admit a Turing instability must exhibit a $+/-$ ecological interaction.*

*Proof.* Assume that two species admit a Turing instability. Then, Eq. (S6) implies that

$$d_1 n_2^* \partial_2 f_2(\{n_j^*\}) + d_2 n_1^* \partial_1 f_1(\{n_j^*\}) > 2\sqrt{d_1 d_2 \text{det } J(0)} \geq 0 \quad \text{[S8]}$$

Since $n_i^*$ is a stable fixed point of the temporal dynamics,

$$0 > \text{tr } J(0) = n_2^* \partial_2 f_2(\{n_j^*\}) + n_1^* \partial_1 f_1(\{n_j^*\}). \quad \text{[S9]}$$

These two equations can only be satisfied if $\partial_2 f_2(\{n_j^*\})$ and $\partial_1 f_1(\{n_j^*\})$ have opposite signs. Moreover, as $n_i^*$ is a stable fixed point of the temporal dynamics,

$$0 < \text{det } J(0) = n_2^* n_1^* [\partial_2 f_2(\{n_j^*\}) \partial_1 f_1(\{n_j^*\}) - \partial_1 f_2(\{n_j^*\}) \partial_2 f_1(\{n_j^*\})]. \quad \text{[S10]}$$

Since $\partial_2 f_2(\{n_j^*\})$ and $\partial_1 f_1(\{n_j^*\})$ have opposite signs, this inequality can only be satisfied if $\partial_1 f_2(\{n_j^*\})$ and $\partial_2 f_1(\{n_j^*\})$ have opposite signs. In particular, the fecundity of one species must increase with the density of the other, while the fecunding of the other must decrease. In particular, the two species must exhibit a $+/-$ ecological interaction. □

This result can be understood intuitively. The predators act as a global inhibitor of patterns (population $i = I$, inhibitor) and their prey acts as a local activator of patterns (population $i = A$, activator), which is described by the inequalities $\partial_I f_A(\{n_j^*\}) < 0$ and $\partial_A f_I(\{n_j^*\}) > 0$. From the proof of Lemma 4, it is also true that $\partial_I f_I(\{n_j^*\})$ and $\partial_A f_A(\{n_j^*\})$ must have opposite signs for patterns to form. Indeed, when the predator abundance is artificially increased from the equilibrium value, there is a lack of prey and the fecundity of predators decreases, suggesting that $\partial_I f_I(\{n_j^*\}) < 0$. In contrast, if the prey abundance is artificially increased from the equilibrium value, the predation per individual prey decreases and the fecundity of prey increases, suggesting that $\partial_A f_A(\{n_j^*\}) > 0$. With these assumptions about partial derivatives, I can prove the main result of this section.

**Theorem 1.** *The spatially homogeneous steady state $n_i(t,x) = n_i^*$ of Eq. (3) is unstable to spatial perturbations (i.e., admits a Turing instability) precisely when*

$$d_A < \alpha d_I, \tag{S11}$$

*for some $\alpha \in [0,1]$ that is independent of the motility $d_i$ of each species $i$.*

*Proof.* The spatially homogeneous steady state $n_i(t,x) = n_i^*$ of the spatiotemporal system in Eq. (3) is unstable to spatial perturbations precisely if there is a Turing or a Turing-Hopf instability. By Lemma 2, Turing-Hopf instability cannot emerge. By Lemma 3, Turing instability emerges precisely when

$$\sqrt{\frac{d_A}{d_I}} n_I^* \partial_I f_I(\{n_j^*\}) + \sqrt{\frac{d_I}{d_A}} n_A^* \partial_A f_A(\{n_j^*\}) > 2\sqrt{\det J(0)}. \tag{S12}$$

Since $\partial_I f_I(\{n_j^*\}) < 0$ and $\partial_A f_A(\{n_j^*\}) > 0$, this condition for spatial instability can be manipulated into a single condition

$$d_A < \alpha d_I \tag{S13}$$

with

$$\alpha = \left( \frac{\sqrt{\det J(0)} - \sqrt{-n_I^* n_A^* \partial_A f_I(\{n_j^*\}) \partial_I f_A(\{n_j^*\})}}{n_I^* \partial_I f_I(\{n_j^*\})} \right)^2. \tag{S14}$$

Clearly, $\alpha \geq 0$. If $\alpha > 1$, the homogeneous steady state would be unstable to spatial perturbations for motilities $d_A = d_I$. But, in this case, the eigenvalues of $J(k^2) = J - d_A k^2 I$ would be the eigenvalues of $J$ shifted by $-d_A k^2$. In particular, their real parts would be negative for any mode $k$ and the homogeneous steady state would be stable. This is a contradiction. Thus, $\alpha \leq 1$. □

### 3. Evolutionary dynamics

To explore the evolution of motility, I assume that mutations can introduce phenotypic heterogeneity into populations of each species. In particular, the spatiotemporal model in Eq. (S2) is generalised to incorporate multiple motility phenotypes $p$ with motility strategies $d_{i,p}$. More specifically, I assume that phenotypes $p$ for each species $i$ make a phenotypic set $\mathcal{P}_i$ and that different phenotypes correspond to different motility strategies, that is $d_{i,p} \neq d_{i,q}$ whenever $p \neq q$. In practice, this condition can be achieved by considering motility values $d_{i,p} = p\Delta d_i$ for phenotypes $p = 0, \ldots, P$, which are separated by small steps in motility $\Delta d_i$. For each phenotype $p$, the population density $n_i(t,x,d_{i,p})$ is governed by Eq. (5)

$$\partial_t n_i(t,x,d_{i,p}) = d_{i,p} \partial_x^2 n_i(t,x,d_{i,p}) + n_i(t,x,d_{i,p}) f_i \left( \left\{ \sum_p n_j(t,x,d_{j,p}) \right\} \right), \tag{S15}$$

with no-flux boundary conditions $\partial_x n_i(t,x,d_{i,p})|_{x=0,L} = 0$.

This section aims to study the properties of steady states to equation Eq. (S15). To start, I define the total fitness of a species by the total fecundity

$$F_i(t) = \int f_i \left( \left\{ \sum_p n_j(t,x,d_{j,p}) \right\} \right) dx, \tag{S16}$$

and explore how it behaves at a steady state of the eco-evolutionary dynamics.

**Theorem 2.** *If $n_j^*(x,d_{j,p})$ is a steady state of the eco-evolutionary model in* Eq. (S15) *and $F_i^*$ the corresponding total fitness of species $i$, then the total fitness is bounded above by $F_i^* \leq 0$ and attains the upper bound precisely when all phenotypes $p$ satisfy $\partial_x n_i^*(x,d_{i,p}) = 0$, i.e., there are no static patterns.*

*Proof.* The steady-state $n_i^*(x,d_{i,p})$ of Eq. (S15) satisfies

$$0 = d_{i,p} \partial_x^2 n_i^*(x,d_{i,p}) + n_i^*(x,d_{i,p}) f_i \left( \left\{ \sum_p n_j^*(x,d_{j,p}) \right\} \right). \tag{S17}$$

Therefore, the total fitness satisfies

$$F_i^* = \int f_i\left(\left\{\sum_p n_j^*(x,d_{j,p})\right\}\right)\,dx = -d_{i,p}\int \frac{\partial_x^2 n_i^*(x,d_{i,p})}{n_i^*(x,d_{i,p})}\,dx = -d_{i,p}\int\left(\frac{\partial_x n_i^*(x,d_{i,p})}{n_i^*(x,d_{i,p})}\right)^2 dx \le 0, \tag{S18}$$

with equality precisely when $\partial_x n_i^*(x,d_{i,p}) = 0$ identically for all phenotypes $p$. I use Eq. (S17) in the first step and integration by parts with no-flux boundary conditions $\partial_x n_i^*(x,d_{i,p})|_{x=0,L} = 0$ in the second step. □

Similarly as steady states of the temporal dynamics in Eq. (S1) become steady states of the spatiotemporal dynamics in Eq. (S2), the steady states of the spatiotemporal dynamics in Eq. (S2) become steady states of the eco-evolutionary dynamics in Eq. (S15).

**Lemma 5.** *Any steady state $n_j^*(x)$ of the system in* Eq. (S2) *with fixed motility $d_j$ defines a steady state $n_j^*(t,x,d_{j,p})$ of the system in* Eq. (S15) *that satisfies*

$$n_j(t,x,d_{j,p}) = \begin{cases} n_j^*(x) & \text{if } d_{j,p} = d_j,\\ 0 & \text{if } d_{j,p} \neq d_j,\end{cases} \tag{S19}$$

*provided the phenotypic sets $\mathcal{P}_j$ of each species $j$ include a phenotype $p$ with motility $d_{j,p} = d_j$.*

*Proof.* Assume that a phenotypic set $\mathcal{P}_j$ of each species $j$ includes a phenotype $p$ with motility $d_{j,p} = d_j$, and consider a function given by

$$n_j(t,x,d_{j,p}) = \begin{cases} n_j(t,x) & \text{if } d_{j,p} = d_j,\\ 0 & \text{if } d_{j,p} \neq d_j.\end{cases} \tag{S20}$$

Then, $n_j(t,x,d_{j,p})$ solves Eq. (S15) precisely if $n_j(t,x)$ solves Eq. (S2). The correspondence between steady states follows. □

Having identified steady states of the spatiotemporal dynamics with steady states of the eco-evolutionary dynamics, the evolutionary stability of these steady states can be examined, giving rise to the main result of this work.

**Theorem 3.** *Assume that $n_j^*(x)$ is a positive, stable and spatially inhomogeneous steady state of* Eq. (S2) *(i.e., a Turing pattern) and consider solutions of* Eq. (S15)

$$\begin{aligned} n_j(t,x,d_j) &= n_j^*(x) + n_j'(t,x) && \text{for both populations } j,\\ n_i(t,x,d) &= m(t,x) && \text{for a mutant in population } i,\end{aligned} \tag{S21}$$

*with a mutant phenotype of species $i$ with motility $d$. Then, the solution $m = 0$, $n_i' = 0$ is locally asymptotically stable (resp. unstable) if $d > d_i$ (resp. $d < d_i$). In particular, the steady state of* Eq. (S15)

$$n_j(t,x,d_{j,p}) = \begin{cases} n_j^*(x) & \text{if } d_{j,p} = d_j,\\ 0 & \text{if } d_{j,p} \neq d_j.\end{cases} \tag{S22}$$

*is locally asymptotically unstable whenever there exists a phenotype $p$ of one of the populations $i$ with motility $d_{i,p} < d_i$.*

*Proof.* Assume that population $i$ consists of phenotypes with motility strategies $d_i$ and $d$, while any other population $j$ consists of a single phenotype with motility strategy $d_j$. Furthermore, consider the Turing pattern $n_j^*(x)$ and the perturbation given by Eq. (S21). From from Eq. (S2), Turing pattern $n_j^*(x)$ must satisfy

$$0 = d_i\partial_x^2 n_i^* + n_i^* f_i(\{n_j^*\}),$$

and the perturbed densities $n_j'(t,x)$ and $m(t,x)$ must satisfy Eq. (S15)

$$\begin{aligned} \partial_t m &= d\partial_x^2 m + m f_i(\{n_k^* + n_k' + \mathbf{1}_{k=i} m\})\\ \partial_t n_j' &= d_j\partial_x^2 n_j' + n_j' f_j(\{n_k^* + n_k' + \mathbf{1}_{k=i} m\}) + n_j^*(f_j(\{n_k^* + n_k' + \mathbf{1}_{k=i} m\}) - f_j(\{n_k^*\})),\end{aligned}$$

If these equations are linearised around $m = 0$, $n_j' = 0$, they reduce to

$$\begin{aligned} \partial_t m &= d\partial_x^2 m + m f_i(\{n_k^*\})\\ \partial_t n_j' &= d_j\partial_x^2 n_j' + n_j' f_j(\{n_k^*\}) + n_j^*\sum_l n_l'\partial_l f_j(\{n_k^*\}) + n_j^* m\partial_i f_j(\{n_k^*\}),\end{aligned}$$

These equations can be solved by $(m(t,x), n_j'(t,x)) = e^{\lambda t}(X(x), X_j(x))$, which leads to an eigenvalue problem of the form

$$\lambda\begin{pmatrix} X\\ X_i\end{pmatrix} = \mathcal{L}\begin{pmatrix} X\\ X_i\end{pmatrix} = \left(\begin{array}{c|c} \mathcal{L}_0(d) & 0\\ \hline n_j^*\partial_i f_j(\{n_l^*\}) & \mathcal{L}_{jk}\end{array}\right)\begin{pmatrix} X\\ X_k\end{pmatrix},$$

where

$$\mathcal{L}_0(d) = d\partial_x^2 + f_i(\{n_k^*\})$$
$$\mathcal{L}_{jj} = d_j\partial_x^2 + f_j(\{n_l^*\}) + n_j^*\partial_j f_j(\{n_l^*\}) \quad \text{[S23]}$$
$$\mathcal{L}_{jk} = n_j^*\partial_k f_j(\{n_l^*\}) \text{ whenever } j \neq k.$$

The eigenvalues of the operator $\mathcal{L}$ determine the stability of the steady state $m = 0$, $n_j' = 0$. This steady state is locally asymptotically unstable (resp. stable) if the operator $\mathcal{L}$ has an eigenvalue with a positive real part (resp. has all eigenvalues with negative real parts). As the operator $\mathcal{L}$ is block lower-diagonal*, the set of its eigenvalues is a union of the sets of eigenvalues for $\mathcal{L}_0(d)$ and $\mathcal{L}_{ij}$. Operator $\mathcal{L}_{ij}$ describes the linearised growth of perturbations to the spatio-temporal system in Eq. (S2) around the Turing pattern $n_j^*(x)$. By assumption, this steady state is stable and all eigenvalues of the operator $\mathcal{L}_{ij}$ have negative real parts. Therefore, the stability of the steady state $m = 0$, $n_j' = 0$ is determined solely by the eigenvalues of $\mathcal{L}_0(d)$, which can be analyzed by modification of ideas from (1). In particular, it can be shown that $\mathcal{L}_0(d)$ has an eigenvalue with a positive real part precisely if $d > d_i$.

To prove this claim, notice that the operator $\mathcal{L}_0(d)$ is self-adjoint and its eigenvalues must be real. Moreover, its eigenvalues and eigenfunctions are all analytic functions of $d$ (2). Now, fix $d = d_i$ and notice that $\mathcal{L}_0(d_i)n_i^*(x) = 0$. So, $\mathcal{L}_0(d_i)$ has 0 as an eigenvalue. Since $n_i^*(x)$ is positive and only the largest eigenvalue can have a positive eigenfunction (3), all other eigenvalues of $\mathcal{L}_0(d_i)$ are negative. The proof of the claim is completed by showing that whenever the largest eigenvalue of the operator $\mathcal{L}_0(d)$ is zero, the derivative of the eigenvalue with respect to $d$ is negative. If $X(x, d)$ is the normalised eigenfunction of the operator $\mathcal{L}_0(d)$, then the corresponding eigenvalue is (4)

$$\int [-d(\partial_x X)^2 + f_i X^2]\, dx.$$

The derivative of this eigenvalue with respect to $d$ is

$$-\int (\partial_x X)^2\, dx + 2\int [-d\partial_x X \partial_d \partial_x X + f_i X \partial_d X]\, dx = -\int (\partial_x X)^2\, dx + 2\int X\partial_d[d\partial_x^2 X + f_i X]\, dx = -\int (\partial_x X)^2\, dx,$$

where the first equality uses integration by parts (with no-flux boundary conditions $\partial_x X = 0$) and the second equality assumes that the eigenvalue of $\mathcal{L}_0(d)$ corresponding to $X$ is 0. Therefore, the claim that this derivative is negative follows if $X$ is not a spatially constant function. This must be true since $d\partial_x^2 X + f_i X = 0$ and $f_i(\{n_j^*(x)\})$ is not identically zero because of the spatial inhomogeneity of the Turing pattern $n_j^*(x)$. This proves the claim and the result of the theorem follows. □

In particular, Theorem 3 shows that motility strategies consistent with the formation of Turing patterns are not evolutionarily stable. Importantly, this result is general and does not depend on the specific choice of fecundity functions $f_i$ or the number of interacting species.

### 4. Spatiotemporal dynamics of two species with heterogeneous motility

In section 2, I showed that two species with fixed motility strategies cannot exhibit dynamic patterns, but can only admit Turing patterns. In section 3, I showed that Turing patterns are evolutionarily unstable if motility can evolve. However, once evolution is introduced into the modelling, it is possible that each species can harbour multiple motility phenotypes. While it is not feasible to analyze the model with arbitrarily many phenotypes analytically, I will extend the analysis of pattern formation to situations where one of the two species harbours two different motility strategies.

I assume that species 1 has motility strategies $d_{1,1}$ and $d_{1,2}$, while species 2 has a fixed motility $d_{2,1} \neq 0$, with population densities evolving according to Eq. (S15). Since the length of the domain $L$ is assumed to be large, I will consider the limiting behaviour of the system when $L \to \infty$. Under these conditions, the spatial variable $x$ can be rescaled as $x \to x/\sqrt{d_{2,1}}$. Defining $d = d_{1,1}/d_{2,1}$ and $d' = d_{1,2}/d_{2,1}$, Eq. (S15) for population densities reduces to

$$\partial_t n_1 = n_1 f_1(n_1 + n_1', n_2) + d\partial_x^2 n_1$$
$$\partial_t n_1' = n_1' f_1(n_1 + n_1', n_2) + d'\partial_x^2 n_1' \quad \text{[S24]}$$
$$\partial_t n_2 = n_2 f_2(n_1 + n_1', n_2) + \partial_x^2 n.$$

This dynamical system admits a line of homogeneous stationary states, parametrised by the relative abundance $q \in [0, 1]$ of the phenotype with motility $d$, and is given by

$$n_1 = qn_1^*$$
$$n_1' = (1 - q)n_1^* \quad \text{[S25]}$$
$$n_2 = n_2^*$$

where $f_i(n_1^*, n_2^*) = 0$. To analyze the stability of these steady states, I first prove the following lemma.

*The claim presented in this sentence is true for operators on finite-dimensional vector spaces, and therefore true for any discretization of the operator. The proof is immediate as a determinant of a block lower-diagonal matrix is a product of determinants of the block matrices on the diagonal, and therefore, the characteristic polynomial factorises into the characteristic polynomials of the diagonal block matrices. For infinite-dimensional vector spaces, this claim requires further investigation that goes beyond the purposes of this paper.

**Lemma 6.** *Let*

$$g(x) = a_3x^3 + a_2x^2 + a_1x + a_0 \tag{S26}$$

*be a cubic function with* $a_3 < 0$ *and* $a_0 \leq 0$, *and let*

$$h(x) = b_2x^2 + b_1x + b_0 \tag{S27}$$

*be a quadratic function with* $b_2 > 0$ *and* $b_0 > 0$. *Then, there is a positive number* $x' > 0$ *such that* $g(x') \geq 0$ *precisely if the following conditions are satisfied*

$$\begin{aligned} &0 < a_2^2 - 3a_1a_3 \\ &0 < a_2 + \sqrt{a_2^2 - 3a_1a_3} \\ &0 < 2a_2^3 + 2(a_2^2 - 3a_1a_3)^{3/2} - 9a_1a_2a_3 + 27a_0a_3^2. \end{aligned} \tag{S28}$$

*Furthermore, there is a positive number* $x' > 0$ *such that* $g(x') \geq 0$ *and* $h(x') > 0$ *precisely if all conditions in* Eq. (S28) *are satisfied and at least one of the following conditions is not*

$$\begin{aligned} b_1 &< -\sqrt{4b_2b_0} \\ 3a_3(b_1 + \sqrt{b_1^2 - 4b_2b_0}) &\leq 2b_0(a_2 + \sqrt{a_2^2 - 3a_1a_3}) \\ 2b_0(a_2 + \sqrt{a_2^2 - 3a_1a_3}) &\leq 3a_3(b_1 - \sqrt{b_1^2 - 4b_2b_0}) \\ g\left(\frac{-b_1 - \sqrt{b_1^2 - 4b_2b_0}}{2b_2}\right) &\leq 0 \\ g\left(\frac{-b_1 + \sqrt{b_1^2 - 4b_2b_0}}{2b_2}\right) &\leq 0. \end{aligned} \tag{S29}$$

*Proof.* Notice that the cubic $g(x)$ has a derivative

$$g'(x) = 3a_3x^2 + 2a_2x + a_1. \tag{S30}$$

This is a quadratic function with a negative leading coefficient, leading to two different cases depending on the existence of solutions to the equation $g'(x) = 0$. If $g'(x) = 0$ does not have any solutions, $g'(x) < 0$ for all $x$ and $g(x)$ is monotone decreasing. Since $g(0) = a_0 \leq 0$, this implies $g(x) < 0$ for all $x > 0$, contradicting the existence of $x'$. In contrast, if $g'(x) = 0$ has solutions, it must be true that

$$0 < a_2^2 - 3a_3a_1 \tag{S31}$$

and the solutions are given by

$$x_\pm = \frac{-a_2 \pm \sqrt{a_2^2 - 3a_1a_3}}{3a_3} \tag{S32}$$

with $x_+ < x_-$. Since $a_3 < 0$, the solution $x_+$ corresponds to a local minimum while $x_-$ corresponds to a local maximum. If $x_- < 0$, then $g(x)$ is monotone decreasing for $x \geq 0$, contradicting the existence of $x'$ by the previous argument. Therefore, for $x'$ to exist, it must be true that $x_- > 0$ or equivalently

$$0 < a_2 + \sqrt{a_2^2 - 3a_1a_3}. \tag{S33}$$

Finally, there is an $x'$ with $g(x') > 0$ precisely if the value of $g(x_-)$ at the maximum $x_-$ is positive. Upon a bit of algebra, this condition is equivalent to

$$0 < 2a_2^3 + 2(a_2^2 - 3a_1a_3)^{3/2} - 9a_1a_2a_3 + 27a_0a_3^2. \tag{S34}$$

If we additionally require that $h(x') > 0$, previous conditions must be satisfied and the region where $g(x) > 0$ must not be included in the region where $h(x) \leq 0$. Since $b_2 > 0$, the region $h(x) \leq 0$ can include the open set given by $g(x) > 0$ only if $h(x)$ has two roots, that is only if

$$b_1 > -\sqrt{4b_2b_0}. \tag{S35}$$

If the quadratic $h(x)$ has two roots

$$x_{1,2} = \frac{-b_1 \pm \sqrt{b_1^2 - 4b_2b_0}}{2b_2} \tag{S36}$$

with $x_1 < x_2$, the region of $g(x) > 0$ is included in the region of $h(x) \leq 0$ precisely if $x_1 \leq x_+ < x_- \leq x_2$ and $g(x_{1,2}) \leq 0$. The conditions in Eq. (S29) is derived by writing down these conditions in detail. □

With this lemma, it is possible to analyze what patterns form in a model for two species with phenotypic heterogeneity.

**Theorem 4.** *The steady state in* Eq. (S25) *with relative abundance $q$ of the phenotype with motility $d$ admits a Turing instability if the coefficients*

$$\begin{aligned} a_3 &= -dd' \\ a_2 &= (d(1-q)+d'q)n_1^*\partial_1 f_1 + dd'n_2^*\partial_2 f_2 \\ a_1 &= -(d(1-q)+d'q)n_1^*n_2^*(\partial_1 f_1\partial_2 f_2 - \partial_1 f_2\partial_2 f_1) \\ a_0 &= 0 \end{aligned} \tag{S37}$$

*satisfy all conditions in* Eq. (S28). *The steady state in* Eq. (S25) *with relative abundance $q$ of the phenotype with motility $d$ admits a Turing-Hopf instability if the coefficients*

$$\begin{aligned} a_3 &= -(d+d')(1+d)(1+d') \\ a_2 &= [(1+d)(1+d') + (d+d')(1+d(1-q)+d'q)]n_1^*\partial_1 f_1 + (d+d')(2+d+d')n_2^*\partial_2 f_2 \\ a_1 &= -2(1+d+d')n_1^*n_2^*\partial_1 f_1\partial_2 f_2 + (1+dq+d'(1-q))n_1^*n_2^*\partial_1 f_2\partial_1 f_2 - (1+d(1-q)+dq)(n_1^*\partial_1 f_1)^2 - (d+d')(n_2^*\partial_2 f_2)^2 \\ a_0 &= (n_1^*\partial_1 f_1 + n_2^*\partial_2 f_2)n_1^*n_2^*(\partial_1 f_1\partial_2 f_2 - \partial_1 f_2\partial_2 f_1) \end{aligned} \tag{S38}$$

*satisfy all conditions in* Eq. (S28) *and the coefficients*

$$\begin{aligned} b_2 &= (d+d'+dd') \\ b_1 &= -(1+d(1-q)+d'q)n_1^*\partial_1 f_1 - (d+d')n_2^*\partial_2 f_2 \\ b_0 &= n_1^*n_2^*(\partial_1 f_1\partial_2 f_2 - \partial_1 f_2\partial_2 f_1) \end{aligned} \tag{S39}$$

*break at least one condition in* Eq. (S29).

*Proof.* The Jacobian of the system in Eq. (S24) is given by

$$J(k^2) = \begin{pmatrix} qn_1^*\partial_1 f_1 - dk^2 & qn_1^*\partial_1 f_1 & qn_1^*\partial_2 f_1 \\ (1-q)n_1^*\partial_1 f_1 & (1-q)n_1^*\partial_1 f_1 - d'k^2 & (1-q)n_1^*\partial_2 f_1 \\ n_2^*\partial_1 f_2 & n_2^*\partial_1 f_2 & n_2^*\partial_2 f_2 - k^2 \end{pmatrix} \tag{S40}$$

The eigenvalues $\lambda$ of this Jacobian satisfy the characteristic equation

$$\lambda^3 - \mathrm{tr}J(k^2)\lambda^2 + \mathrm{tr}[\mathrm{adj}J(k^2)]\lambda - \det J(k^2) = 0, \tag{S41}$$

where

$$\begin{aligned} \mathrm{tr}J(k^2) &= -k^2(d+d'+1) + n_1^*\partial_1 f_1 + n_2^*\partial_2 f_2 \\ \mathrm{tr}[\mathrm{adj}J(k^2)] &= k^4(d+d'+dd') - k^2[(1+d(1-q)+d'q)n_1^*\partial_1 f_1 + (d+d')n_2^*\partial_2 f_2] + n_1^*n_2^*(\partial_1 f_1\partial_2 f_2 - \partial_1 f_2\partial_2 f_1) \\ \det J(k^2) &= -k^6dd' + k^4[(d(1-q)+d'q)n_1^*\partial_1 f_1 + dd'n_2^*\partial_2 f_2] - k^2(d(1-q)+d'q)n_1^*n_2^*(\partial_1 f_1\partial_2 f_2 - \partial_1 f_2\partial_2 f_1). \end{aligned} \tag{S42}$$

By the fundamental theorem of algebra, there are three eigenvalues $\lambda$. Moreover, their sum is negative because

$$\mathrm{tr}J(k^2) = -k^2(d+d'+1) + \mathrm{tr}J(0) < 0. \tag{S43}$$

Therefore, not all eigenvalues can have positive real parts.

If there is a Turing instability at wavelength $k$, a real eigenvalue passes through the line Re $\lambda(k^2) = 0$. Since not all eigenvalues can have positive real parts, either both remaining eigenvalues have negative real parts or one is real positive and the other real negative. In the latter case, there must exist $k' < k$ such that the real positive eigenvalue passes through the line Re $\lambda(k^2) = 0$ and the other two eigenvalues have negative real parts. In summary, Turing instability occurs precisely if a single real eigenvalue passes through the line Re $\lambda(k^2) = 0$, while the remaining two eigenvalues have negative real parts. In particular, as $k$ is varied, the eigenvalues change from all having negative real parts to one being positive real. By the Routh–Hurwitz criterion, all eigenvalues have negative real parts precisely if $\mathrm{tr}J(k^2) < 0$, $\mathrm{tr}[\mathrm{adj}J(k^2)] > 0$, $\det J(k^2) < 0$, and $\mathrm{tr}J(k^2)\mathrm{tr}[\mathrm{adj}J(k^2)] - \det J(k^2) < 0$. Moreover, at the critical wavelength $k$, one eigenvalue passes through 0, implying that $\det J(k^2)$ passes through 0. Since $\det J(k^2) < 0$ if all eigenvalues have negative real parts, a Turing instability occurs precisely if there is a positive $k^2 = x > 0$ such that $\det J(x) > 0$. Application of Lemma 6 to $g(x) = \det J(x)$ implies the result.

If there is a Turing-Hopf bifurcation at wavelength $k$, a pair of complex eigenvalues passes through the line $i\mathbb{R} \setminus \{0\}$, while the remaining eigenvalue is negative real. When this happens, $\lambda(k^2) = i\omega \in i\mathbb{R} \setminus \{0\}$ solves the characteristic equation, that is

$$\begin{aligned} \omega^2\mathrm{tr}J(k^2) &= \det J(k^2) \\ \omega^2 &= \mathrm{tr}[\mathrm{adj}J(k^2)], \end{aligned} \tag{S44}$$

or equivalently

$$\begin{aligned} \mathrm{tr}[\mathrm{adj}J(k^2)]\mathrm{tr}J(k^2) &= \det J(k^2) \\ 0 &< \mathrm{tr}[\mathrm{adj}J(k^2)]. \end{aligned} \tag{S45}$$

Since $\mathrm{tr}J(k^2)\mathrm{tr}[\mathrm{adj}J(k^2)] - \det J(k^2) < 0$ when all eigenvalues have negative real parts, the Turing-Hopf instability appears precisely if there is a positive $k^2 = x > 0$ such that

$$\begin{aligned} 0 &< \mathrm{tr}[\mathrm{adj}J(x)]\mathrm{tr}J(x) - \det J(x) \\ 0 &< \mathrm{tr}[\mathrm{adj}J(x)]. \end{aligned} \quad \text{[S46]}$$

Application of Lemma 6 to $h(x) = \mathrm{tr}[\mathrm{adj}J(x)]$ and

$$\begin{aligned} g(x) = {} & \mathrm{tr}[\mathrm{adj}J(x)]\mathrm{tr}J(x) - \det J(x) = \\ & - x^3(d + d')(1 + d)(1 + d') \\ & + x^2\{[(1 + d)(1 + d') + (d + d')(1 + d(1 - q) + d'q)]n_1^*\partial_1 f_1 + (d + d')(2 + d + d')n_2^*\partial_2 f_2\} \\ & - x[2(1 + d + d')n_1^*n_2^*\partial_1 f_1\partial_2 f_2 - (1 + dq + d'(1 - q))n_1^*n_2^*\partial_1 f_2\partial_1 f_2 + (1 + d(1 - q) + dq)(n_1^*\partial_1 f_1)^2 + (d + d')(n_2^*\partial_2 f_2)^2] \\ & + (n_1^*\partial_1 f_1 + n_2^*\partial_2 f_2)n_1^*n_2^*(\partial_1 f_1\partial_2 f_2 - \partial_1 f_2\partial_2 f_1), \end{aligned} \quad \text{[S47]}$$

implies the result. □

While it is analytically unfeasible to explore how motility affects pattern formation from the set of conditions in Theorem 4, it is possible to plot these conditions numerically (Fig. S1). If prey has two motility strategies and predators have a single motility strategy, the result is similar to the case with prey having a single motility strategy. Specifically, either homogeneous distribution is stable or Turing patterns form (Fig. S1a-c). However, if predators have two motility strategies and prey have a single motility strategy, the linear stability analysis suggests that dynamic patterns can form via the Turing-Hopf bifurcation (Fig. S1d-f).

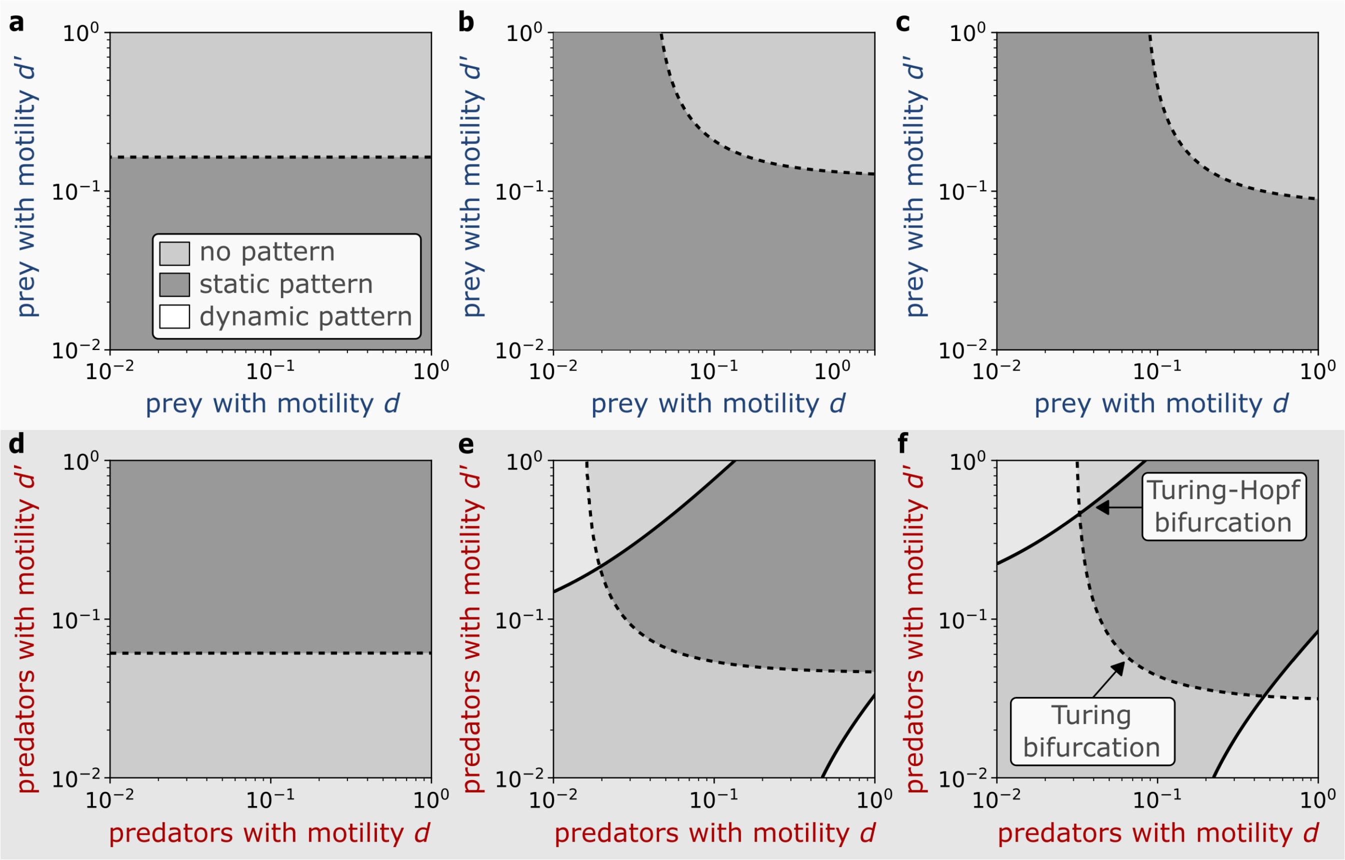


**Fig. S1.** Heterogeneous motility can introduce dynamic patterns. Linear stability analysis of Theorem 4 has been used to plot regions where static (dark grey) and dynamic (white) patterns form. The dashed line corresponds to the onset of static patterns via Turing bifurcation, while the solid line denotes the onset of dynamic patterns via Turing-Hopf bifurcation. (a-c) Heterogeneity in prey motility. The motility of predators is fixed ($d_I = 1$), while prey admits two phenotypes with motility $d$ and $d'$. The initial spatially homogeneous population is assumed to include the phenotype of motility $d$ at a relative abundance $q = 0$ (a), $q = 0.25$ (b) and $q = 0.5$ (c). (d-f) Heterogeneity in predator motility. The motility of prey is fixed ($d_A = 0.01$), while predators admit two phenotypes with motility $d$ and $d'$. The initial spatially homogeneous population is assumed to include the phenotype of motility $d$ at a relative abundance $q = 0$ (d), $q = 0.25$ (e) and $q = 0.5$ (f).

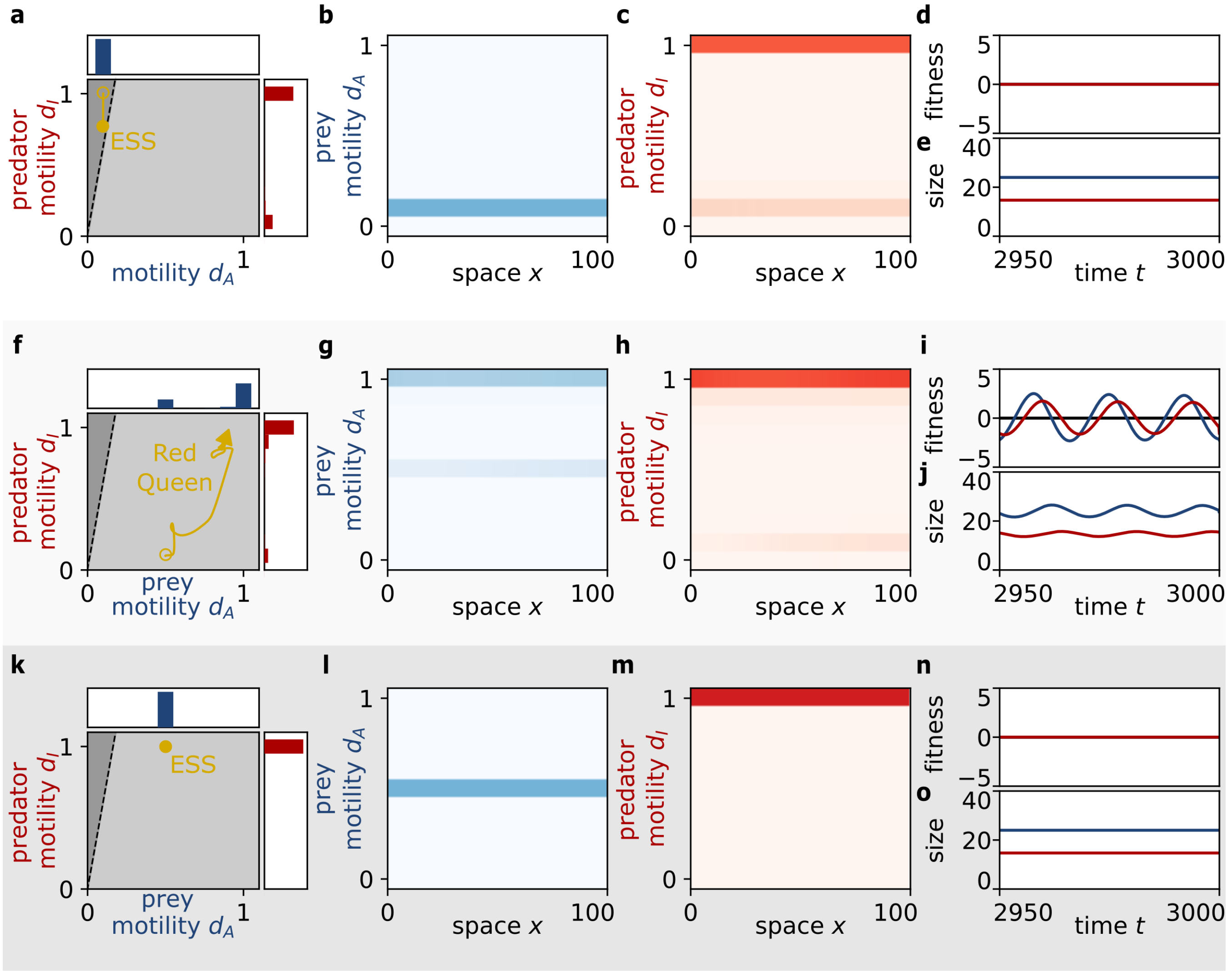


**Fig. S2.** Evolution of motility with mutations of large effect. Analogous to Figure 4, but mutations of large effect are considered (see Methods). (a-d) Stationary evolution, no pattern. The motility of predators (red) and prey (blue) reaches an evolutionarily stable strategy (ESS, golden line) that consists of two predator phenotypes. At this evolutionary equilibrium, there are no patterns in prey density (b) and predator density (c). The total fitness of both species is optimal (d) and the total population size (e) remains constant. (f-j) Dynamic evolution, dynamic pattern. The predators and prey evolve increasing motility (f), leading to predator-prey waves (g,h), and oscillations in total fitness (i) and total population size (j). (k-o) Stationary evolution, no pattern. The motility stays at its original value (k), and the prey (l) and their predators (m) are homogeneously distributed. The total fitness stays at the optimal value (n) and the population size does not change (o).

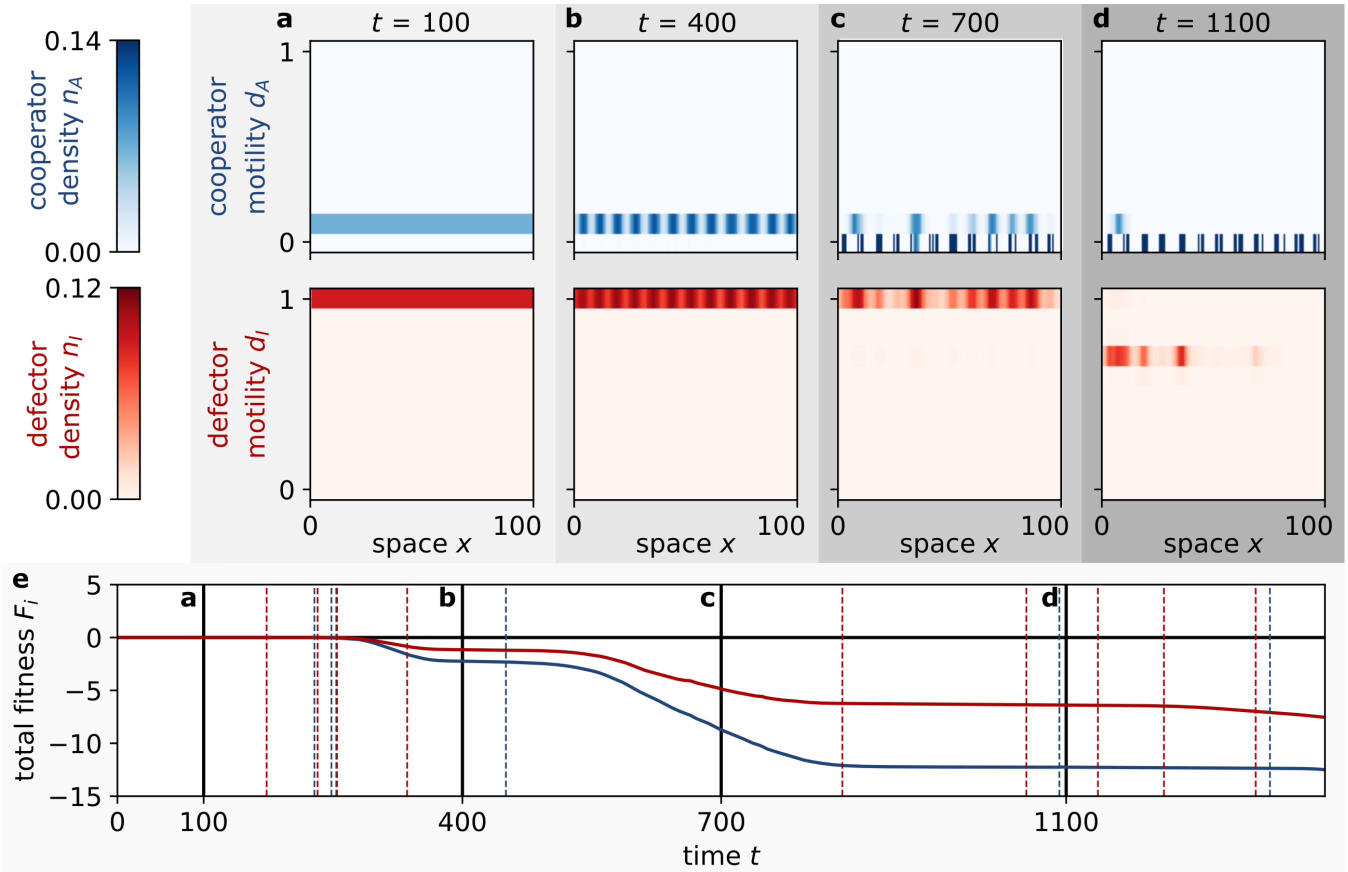


**Fig. S3.** Turing patterns negatively impact the fitness and motility of cooperators and defectors. Analogous to Figure 2, with cooperators and defectors considered instead of predators and prey (see Methods). (a-d) Snapshots. Cooperators (blue) and predators (red) have phenotypes of different motility. Initially, both populations have a single motility phenotype and are nearly spatially homogeneous (a). For appropriate initial motility, cooperators and defectors self-organise to form Turing patterns (b). At a stochastic rate $\mu_i$, phenotypes of each species $i$ are updated by natural selection and mutations. Consequently, the cooperators decrease their motility (c) and so do the defectors (d). In turn, changes in motility trigger changes in the Turing pattern. (e) Evolution of total fitness. The total fitness of cooperators (blue) and defectors (red) is defined by the total fecundity of each population. When Turing patterns form, the total fitness decreases. Vertical black lines indicate the time of snapshots in panels (a-d), horizontal black line corresponds to the optimal total fitness, and dashed lines correspond to the times when phenotypes of cooperators (blue) and defectors (red) are updated.

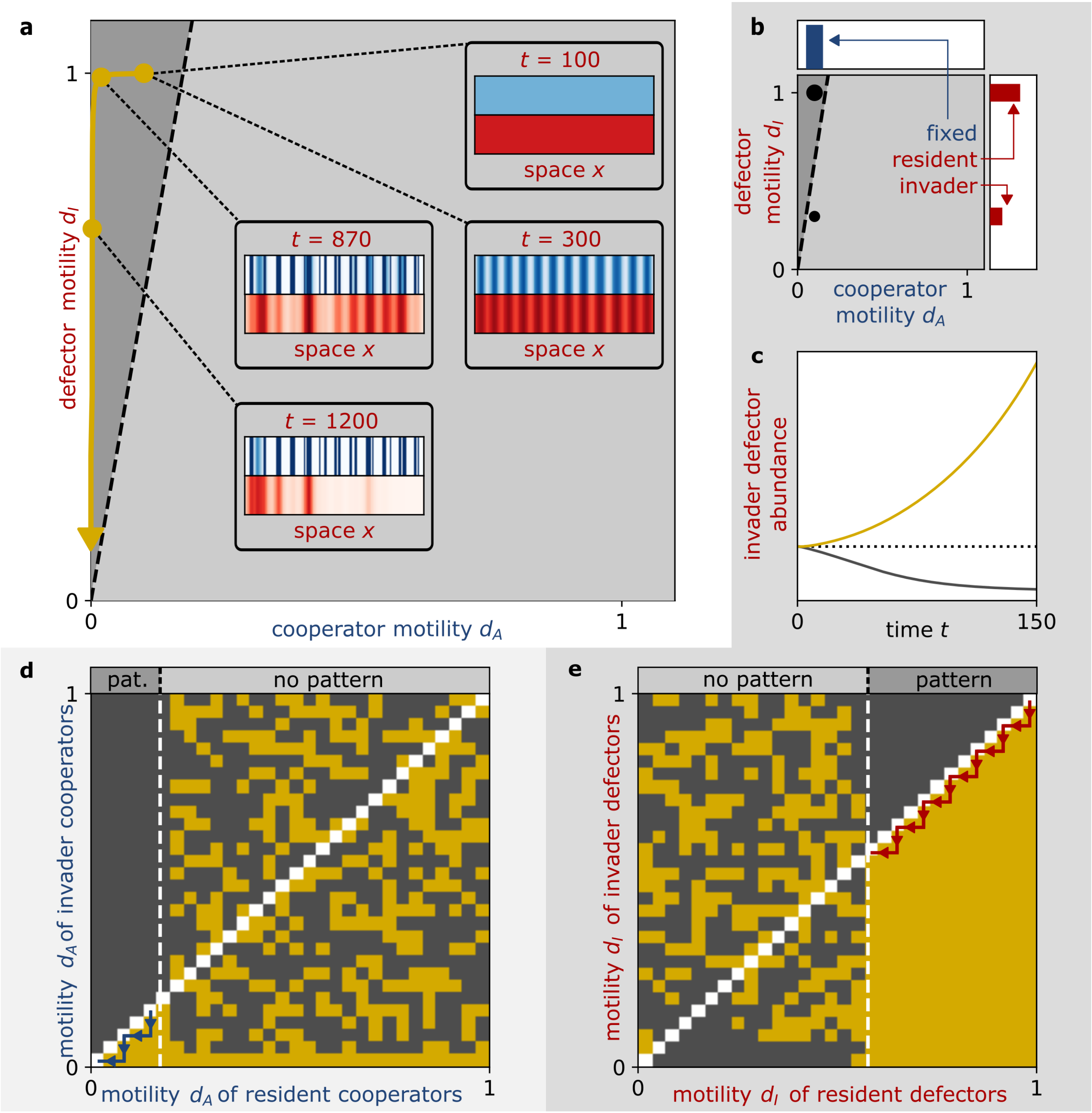


**Fig. S4.** Evolution selects phenotypes of smaller motility when cooperators and defectors form Turing patterns. Analogous to Figure 3, with cooperators and defectors considered instead of predators and prey (see Methods). (a) Evolution of motility. The evolution is indicated by the trajectory of mean coopeartor and defector motility (golden line). The initial motility is compatible with Turing patterns (dark grey). Once patterns form, cooperators with smaller motility are naturally selected, followed by defectors of smaller motility being selected. This leads to serial changes in the spatial patterns of cooperators (blue) and defectors (red), illustrated by snapshots of the total spatial density. (b-c) Invasibility analysis. The cooperator motility is fixed and defectors with resident motility have reached a stable ecological equilibrium with cooperators (b). Subsequently, defectors with invader motility are introduced into the system at low density (b) and the population dynamics is simulated until time $1/\mu_I$, where $\mu_I$ is the defector mutation rate (c). If the defector with invader motility increases in abundance above its initial density (dotted line), the invader phenotype can invade the resident phenotype (yellow), otherwise not (grey) (c). (d) Pairwise invasibility plot for cooperator motility. The defector motility is fixed ($d_I = 1$) and the cooperator motility ($d_A$) is varied. The white dashed line denotes the threshold for the emergence of Turing patterns in the resident population ($d_A < \alpha d_I$). If Turing patterns exist, smaller motility always invades (blue arrows). When Turing patterns are absent, the pairwise invasibility analysis is inconclusive and evolution is neutral. (e) Pairwise invasibility plot for defector motility. Same as (d), but the cooperator motility is fixed ($d_A = 0.1$) and the defector motility ($d_I$) is varied. Again, if Turing patterns exist, defectors with smaller motility invade (red arrows).

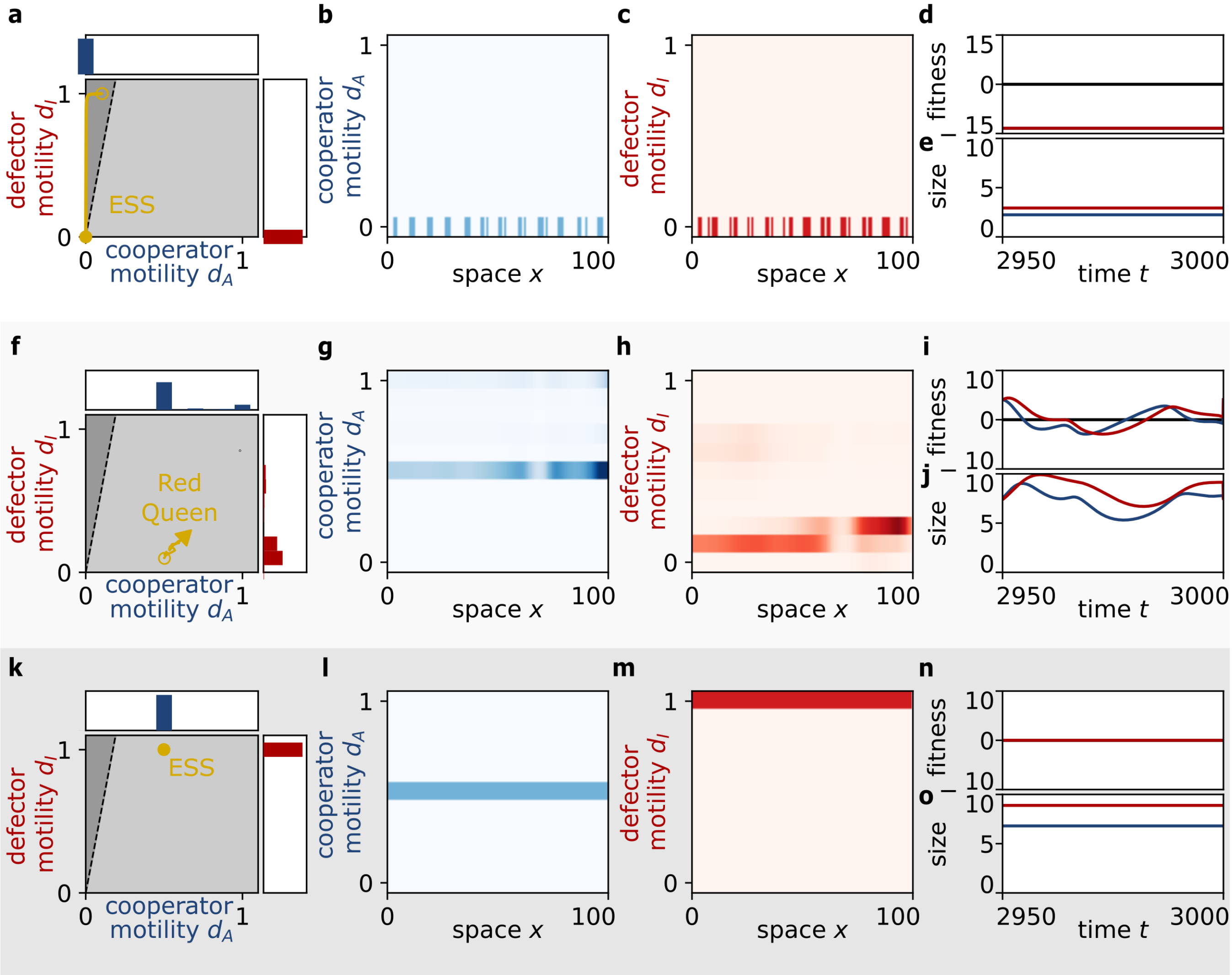


**Fig. S5.** Evolution of motility can introduce complex spatiotemporal patterns of cooperators and defectors. Analogous to Figure 4, with cooperators and defectors considered instead of predators and prey (see Methods). (a-d) Stationary evolution, imprinted pattern. At evolutionary equilibrium (ESS, golden line), cooperators (blue) and defectors (red) lose their motility (a). The patterns of cooperators (b) and defectors (c) along the evolutionary trajectory become imprinted into the equilibrium spatial distribution. Peaks of the transient Turing patterns become regions of coexistence, while valleys become regions of local extinction. The regions of extinction decrease the total fitness of both populations (d) and the population size remains constant (e). (f-j) Dynamic evolution, dynamic pattern. The cooperators and defectors evolve increasing motility (f), leading to chaotic waves (g,h), oscillations in total fitness (i) and total population size (j). (k-o) Stationary evolution, no pattern. The motility stays at its original value (k), and the cooperators (l) and defectors (m) are homogeneously distributed. The total fitness stays at the optimal value (n) and the population size does not change (o).

**Movie S1. Predators and prey of motility consistent with Turing patterns. The spatial and phenotypic distribution of predators (red, lower left panel) and their prey (blue, upper left panel) is plotted at each time step. The phenotypic structure of the populations is represented on the predator-prey motility plane (upper right panel), to show if Turing patterns can form (dark grey background) or not (light grey background). The total fitness of the predator and prey populations (lower right panel, solid line) is compared with the optimal value (black line). The total population size is plotted with dotted lines in the same panel. Mutations are indicated with vertical dashed lines. Parameters were chosen the same as for Figs. 2, 3a and 4a-e.**

**Movie S2. Predators and prey of motility inconsistent with Turing patterns. See the legend of Movie 1 for details. Parameters were chosen the same as for Fig. S24a-e.**

**Movie S3. Predators and prey of motility consistent with Turing patterns subjected to large-effect mutations. Dynamic predator-prey waves promote the evolution of motility. See the legend of Movie 1 for details. Parameters were chosen the same as for Fig. 4f-j.**

**Movie S4. Cooperators and defectors of motility consistent with Turing patterns. See the legend of Movie 1 for details, with cooperators playing the role of prey and defectors playing the role of predators. Parameters were chosen the same as for Figs. S3, S4a and S5a-e.**
